# Individual variation in brain structural-cognition relationships in aging

**DOI:** 10.1101/2021.02.19.431732

**Authors:** Raihaan Patel, Clare E. Mackay, Michelle G. Jansen, Gabriel A. Devenyi, M. Clare O’Donoghue, Mika Kivimäki, Archana Singh-Manoux, Enikő Zsoldos, Klaus P. Ebmeier, M. Mallar Chakravarty, Sana Suri

## Abstract

The sources of inter- and intra-individual variability in age-related cognitive decline remain poorly understood. We examined the association between 20-year trajectories of cognitive decline and multimodal brain structure and morphology in older age. We used the Whitehall II Study, an extensively characterised cohort with 3T brain magnetic resonance images acquired at older age (mean age = 69.52± 4.9) and 5 repeated cognitive performance assessments between mid-life (mean age = 53.2 ±4.9 years) and late-life (mean age = 67.7 ±4.9). Using non-negative matrix factorization, we identified 10 brain components integrating cortical thickness, surface area, fractional anisotropy, and mean and radial diffusivities. We observed two latent variables describing distinct brain-cognition associations. The first describes variations in 5 structural components associated with low mid-life performance across multiple cognitive domains, decline in reasoning, but maintenance of fluency abilities. The second describes variations in 6 structural components associated with low mid-life performance in fluency and memory, but retention of multiple abilities. Expression of latent variables predicts future cognition 3.2 years later (mean age = 70.87 ±4.9). This data-driven approach highlights brain-cognition relationships wherein individuals degrees of cognitive decline *and* maintenance across diverse cognitive functions that are both positively and negatively associated with cortical structure.

## 1.0 Introduction

Cognitive decline is a well-established aspect of the aging process. Age-related impairments which impact everyday functioning have been reported across a range of cognitive domains (Salthouse, 2010;Tucker-Drob, 2011b). However, significant inter-individual variability has also been observed across cognitive domains including memory, spatial functioning, processing speed and reasoning. While some individuals experience accelerated rates of deterioration, others experience a relative maintenance of cognitive functioning into old age (Pudas et al., 2013; Tucker-Drob & Salthouse, 2011; Wilson et al.,2002). Even within the same individual, some cognitive domains may remain intact whereas other domains are more vulnerable to decline (Tucker-Drob, 2011a). It is unclear whether this intra- and inter-individual variation arises from underlying changes already present in early-to-mid life, or if they are established in older age. Improving our understanding of the sources of this variability is an important step in understanding aging-related changes in cognition.

Numerous magnetic resonance imaging (MRI) studies have suggested that inter-individual variation in cognitive function may be partially explained by differences in brain structure, be it through differences in neurobiology at a given time (brain reserve) or in preservation of brain morphology changes during ageing over time (brain maintenance), (Stern et al., 2020). MRI provides macro- and microstructural measures such as volume, thickness, surface area, diffusivity and fractional anisotropy, each of which convey complementary information about the local morphology, axonal density, organization and myelination of the cerebral cortex (Lerch et al., 2017; Tardif et al., 2016). In healthy aging, these techniques have demonstrated widespread age-related degeneration of brain structure, namely decreases in overall brain volume (Hedman et al., 2012), cortical thickness (Fjell, McEvoy, et al., 2014; Lowe et al.,2019; Shaw et al., 2016), and fractional anisotropy as well as increases in diffusivity (Bartzokis, 2004;Lebel et al., 2012; Marner et al., 2003; Tardif et al., 2016). They have also provided evidence for a brain-cognition link in aging, for example by previously established associations between episodic memory performance with volumes and diffusivity of the medial temporal lobe (Philippi et al., 2016; Raz & Rodrigue, 2006; Reas et al., 2018; Schneider et al., 2019), and between decline in executive functioning and widespread grey matter atrophy (Schneider et al., 2019).

However, most studies to date have considered these micro- and macro-structural MRI metrics individually, without considering the complementary information multiple metrics provide on brain structure, or their potential overlap and interdependencies. Moreover, most studies have considered a priori definitions of cognitive decline. Previous approaches investigating brain-cognition relationships have defined subject groupings based on cognitive trajectories and then assessed group differences in brain structure (Persson et al., 2006; Schneider et al., 2019). For example, individuals have been broadly categorised as cognitive “maintainers”, or “decliners” (Josefsson et al., 2012). A common strategy is to categorize the most severe decliners and compare them to the rest of a cohort (Schneider et al., 2019). This approach involves arbitrary cut-offs, and may be biased by the extremes of the decliner-maintenance dimension, neglecting individuals demonstrating neither sharp decline nor strong maintenance (Persson et al., 2012). Furthermore, broad categorisations of maintainers/decliners may also ignore intra-individual heterogeneity and the differential impact of age across cognitive domains (Goh et al., 2012; Park & Reuter-Lorenz, 2009; Rönnlund et al., 2005).

This study builds on previous work in two important ways. First, we integrate multiple MRI based indices of cortical macro- and microstructure. Incorporating data from multiple MRI modalities is useful as each conveys complementary information, such that the resulting multimodal assessments query a wider range of biological phenomena (Tardif et al., 2016). This approach has enabled fine grained assessments of the cerebral cortex. Glasser et al., for example, incorporated data from structural and functional MRI to delineate a novel parcellation of the cortex, including the identification of de novo areas distinguishable as a result of this strategy (Glasser et al., 2016). Seidlitz et al. integrated multimodal MRI indices of cortical structure to identify morphometric networks, such that areas of the cortex displaying morphometric similarity shared cytoarchitectonic and transcriptional features (Seidlitz et al., 2018). These findings demonstrate the specificity of associations demonstrated using multimodal MRI. Modelling shared covariance across MRI metrics, as opposed to separately analysing each piece of information, allows for a more comprehensive assessment of differences across subjects (Groves et al., 2012). To this end, we expand previous work from our group on multi-modal data-driven parcellation of the hippocampus using non-negative matrix factorization (NMF), and extend this methodology to facilitate integration of multiple modalities into a single analytical framework (R. Patel et al., 2020). In this study we consider five MRI metrics. From structural MRI we derive measurements of cortical thickness (CT), surface area (SA), which have been routinely used to track age-related alterations of brain structure (Dickerson et al., 2009; Fjell, Westlye, et al., 2014; Frangou et al., 2022; Habeck et al., 2020; Lemaitre et al., 2012; Lerch et al., 2017; Querbes et al., 2009; Raz & Rodrigue, 2006; Salat et al., 2004; Storsve et al.,2014; Tamnes et al., 2013). We also incorporated diffusion tensor imaging (DTI) indices of mean and radial diffusivity (MD, RD) and fractional anisotropy (FA). While more commonly associated with the study of fiber structure and organization in brain white matter (Alexander et al., 2007; Assaf, 2019), a number of recent studies have demonstrated their sensitivity to microstructural properties of cortical grey matter (Aggarwal et al., 2015; Assaf, 2019; Douaud et al., 2013; Grydeland et al., 2013; Kleinnijenhuis et al., 2015; Kochunov et al., 2011; P. Lee et al., 2020; McKavanagh et al., 2019; Preziosa et al., 2019; Scola et al., 2010; Seidlitz et al., 2018; Torso, Bozzali, et al., 2021; Torso, Ridgway, et al., 2021; Truong et al.,2014). Though their interpretation in cortical grey matter is complex and still under study, DTI indices have been shown to be sensitive to a range of neurobiological features including myelinated and unmyelinated axons, dendrites, and cell bodies (Edwards et al., 2018). Further, while interrelated to some degree, these measures have each been shown to query unique aspects of microstructure (Uddin et al.,2019), making them complementary in nature. Thus, applied to these five MRI metrics, NMF highlights regions of the brain in which shared patterns of variation occur across a range of macro- and microstructural features (D. D. Lee & Seung, 1999; R. Patel et al., 2020; Robert et al., 2022; Sotiras et al.,2015).

Second, we explore individual differences in brain-cognition relationships without *a priori* designations of cognitive trajectories. We instead use a data-driven approach to probe the relationships between brain structure and cognition. We employ partial least squares (PLS), a multivariate technique used to relate two sets of variables together (McIntosh & Lobaugh, 2004), to identify covarying relationships between brain structure and cognitive performance in a data-driven fashion. This approach sidesteps the need for *a priori* definitions and enables identification of dimensions along which subject-specific brain-cognition relationships exist, instead of distinct decline/maintenance categorizations. Importantly, we then assess biological significance of the identified patterns by exploring how individual variation in these dimensions predicts future cognitive performance.

In this study we analyse data from the Whitehall II cohort, a unique and comprehensive longitudinal dataset which enables study of the relationship between mid- and late-life features. This ongoing study was established in 1985 at University College London and initially included 10,308 British civil servants (Marmot & Brunner, 2005). Longitudinal follow-up occurred at multiple timepoints (defined throughout as Waves). We analyse data collected between 1997 and 2016, collected roughly every five years at Wave 5 (1997-1999), 7 (2002-2004), 9 (2007-2009), 11 (2012-2013), and 12 (2015-2016). At each Wave, information on social, cognitive, and biological data was collected, resulting in a unique source of information to study aging. Eight hundred individuals from Wave 11 were randomly selected to participate in the Whitehall II Imaging sub-study (Imaging Wave, 2012-16) in which structural, diffusion, and functional MRI was collected (Filippini et al., 2014). In this work, our goal is to probe sources of individual variability in the link between cognitive decline and cortical structure. We do this by leveraging the unique Whitehall II Imaging sub-study dataset with multivariate techniques to identify brain-cognition modes of covariance between multimodal MRI measures of late-life brain structure and cognitive trajectories from mid- to late-life. We then examine the biological significance of the brain-cognition relationships by testing their association with future cognitive performance.

## 2.0 Methods

Using the comprehensive lifespan data from the Whitehall II Imaging sub-study (Figure 1A), we analyse longitudinal cognitive trajectories across multiple domains and assess their relationship with late-life cortical structure using surface area (SA), cortical thickness (CT), mean diffusivity (MD), fractional anisotropy (FA) and radial diffusivity (RD). Across multiple structural MRI indices, we model shared covariance using NMF. We use linear mixed effects modeling to extract subject specific indices of baseline performance and change in performance across multiple cognitive tests during a period of 20 years. We then use partial least squares (PLS) to identify distinct patterns of covariance between structure and longitudinal cognitive performance, which we term brain-cognition latent variables (Figure 1B) (McIntosh & Lobaugh, 2004; McIntosh & Mišić, 2013). Finally, we use each individual’s expression of the identified latent variables to predict cognitive function at a follow-up timepoint, approximately 3.2 years later (Wave 12) (Figure 1C).

**Figure 1.**
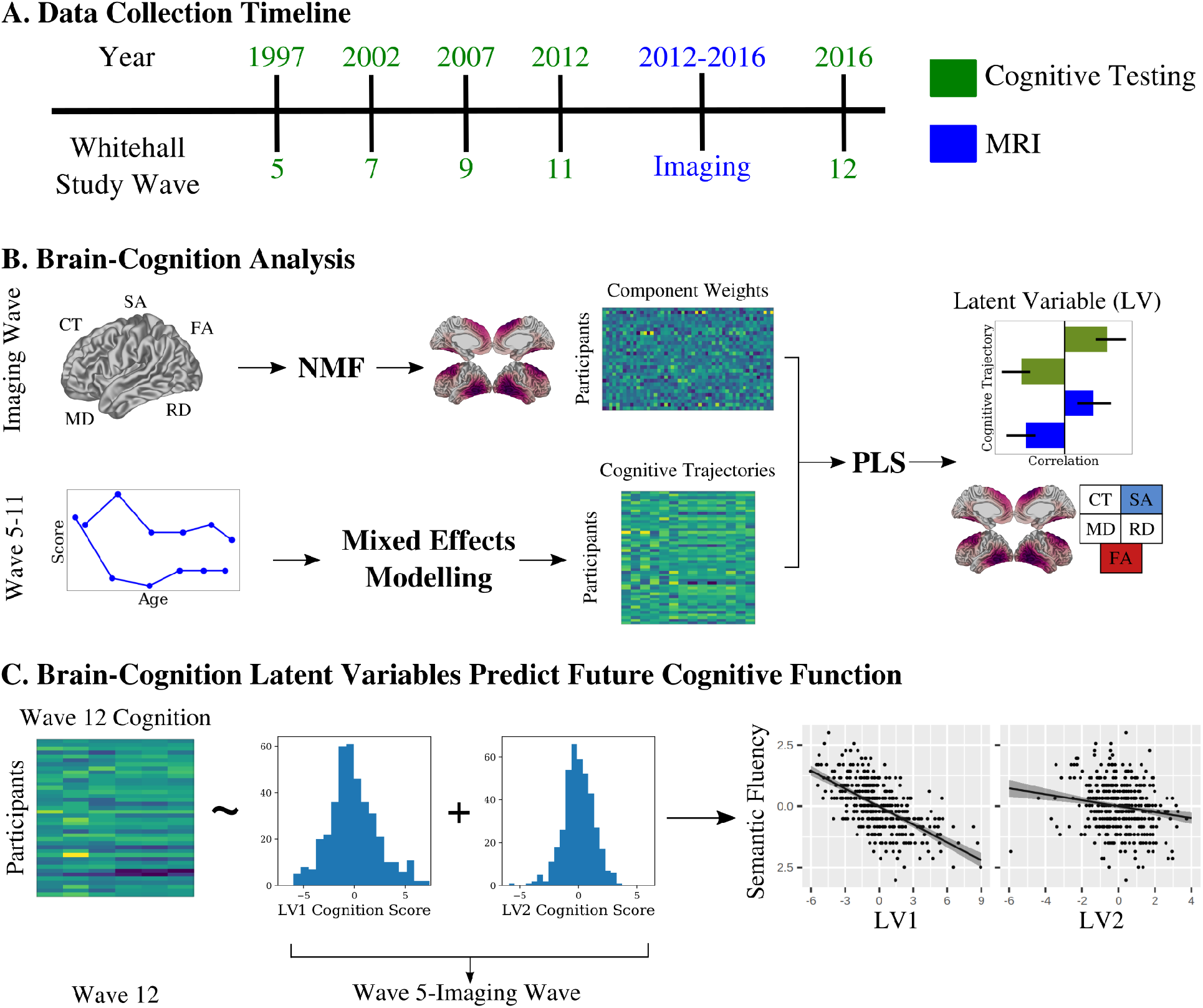
Identifying structural-cognition relationships in aging using data-driven techniques. A) We analysed data from the Whitehall II Imaging Sub-Study. Participants were tested across multiple cognitive domains at ~5-year intervals since 1997, with structural and diffusion MRI collected between Waves 11 and 12. B) We applied non-negative matrix factorization (NMF) to five metrics of cortical morphology: cortical thickness (CT), surface area (SA), fractional anisotropy (FA), and mean and radial diffusivity (MD, RD) to identify patterns of variance. Using cognitive data from Wave 5-11, we applied mixed effects modelling to identify intercept and slope measurements for each individual across a range of cognitive tests during the 20-year period. We then used a partial least squares (PLS) analysis to identify brain-cognition latent variables describing covariance between longitudinal cognitive trajectories and late life cortical morphology. C) We related expression of the brain-cognition latent variables to future cognition to show the identified brain-cognition latent variables predict future performance in two distinct ways.

### 2.1 Sample

We used data from the Whitehall II Imaging Sub-Study (Filippini et al., 2014), a random sample of 800 individuals from the Whitehall II Study of British civil servants, of which 775 received an MRI scan (Marmot & Brunner, 2005). After quality control and exclusions based on available data, 398 participants (mean age = 69.5 years ± 4.2, 92 females) were included in the final sample (Figure S1). These individuals have been assessed longitudinally since 1985 across a total of 12 waves thus far. Cognitive performance was assessed at 5 timepoints at University College London: Wave 5 (1997-1999), Wave 7 (2002-2004), Wave 9 (2007-2009), Wave 11 (2012-2013), and Wave 12 (2015-2016). Structural and diffusion weighted magnetic resonance imaging (MRI) was conducted at the University of Oxford between 2012-2016 (Imaging Wave, Figure 1). Montreal Cognitive Assessment (MOCA) was also conducted at the Imaging Wave to be used as a covariate during subsequent analyses. All participants provided informed written consent, and the Whitehall II Imaging Sub-Study was granted ethical approval by the University of Oxford Central University Research Ethics Committee.

### 2.2 MRI Acquisition

MRI data was acquired on one of two scanners - a 3T Siemens Magnetom Verio (Erlangen, Germany) (n=552) or a 3T Siemens Magnetom Prisma scanner (Erlangen, Germany) (n=223) at the FMRIB Centre in the Wellcome Centre for Integrative Neuroimaging (WIN), Oxford. T1 weighted images were acquired using a Multi-Echo MPRAGE (MEMPR) sequence (1mm^3^, TR = 2530ms, TE = 1.79/3.65/5.51/7.37ms) on the Verio scanner and a closely-matched MPRAGE sequence on the Prisma scanner (1mm^3^, TR=1900ms, TE=3.97ms). Diffusion weighted imaging (DWI) was acquired with an identical sequence across both scanners, using monopolar diffusion encoding gradients with parallel imaging at 2mm isotropic (60 directions, b=1500s/mm^2^, 5 b=0s/mm^2^ images). Detailed acquisition descriptions have been described elsewhere (Filippini et al., 2014).

### 2.3 Obtaining Brain Structural Metrics

In this study we focus on vertex-wise measures of cortical macro- and microstructure. T1w images were preprocessed using the minc-bpipe-library (https://github.com/CoBrALab/minc-bpipe-library), including bias field correction (Tustison et al., 2010), adaptive non-local means denoising (Manjón et al., 2010), head masking and brain extraction (Eskildsen et al., 2012) The resulting bias field corrected, head-masked images and brain masks of each subject were input into the CIVET algorithm (Ad-Dab’bagh et al., 2006;Lerch & Evans, 2005) (version 2.1.0) in order to obtain cortical mid-surfaces and vertex wise measures of cortical thickness (CT) and surface area (SA), describing CT and SA estimates at a total of 81924 points across the cortical mid-surface. Vertex wise CT and SA were blurred using 30 mm and 40 mm geodesic surface kernels, respectively. We masked out 4802 vertices located along the left and right midline as CT and SA estimates in this region are unreliable or nonexistent, resulting in a total of 77122 vertices valid for analysis. CIVET outputs were quality controlled for registration quality, grey/white matter classification accuracy, and surface abnormalities by one of the authors (RP).

DWI data were preprocessed using the FMRIB’s diffusion toolbox (FDT) in order to correct for distortions due to susceptibility, eddy currents, and head motion simultaneously. This process, based on methods applied to the Human Connectome Project Dataset (Sotiropoulos et al., 2013), begins with the topup tool in which pairs of reversed phase encoded images are used to estimate a susceptibility distortion field map (J. L. R. Andersson et al., 2003; Filippini et al., 2014; Sotiropoulos et al., 2013). Next, the eddy tool is used to estimate eddy current distortion as well as head motion using diffusion data acquired with opposite gradient directions (Sotiropoulos et al., 2013). Notably, all distortion estimates are corrected in a single resampling step in order to minimize interpolation and introduction of error (J. Andersson et al.,2012; J. L. R. Andersson et al., 2003; Sotiropoulos et al., 2013). DTIFit (https://fsl.fmrib.ox.ac.uk/fsl/fslwiki/FDT) was used to generate maps of MD, FA, and RD for each subject. For each subject, MD, FA, and RD images were registered to their T1w image using a multispectral affine registration, conducted using the antsRegistration tool from the ANTs toolbox (version 2.3.1) with transform type set to Affine, T1w image as the fixed image, FA, MD, RD images as the moving images. A brain mask for the T1w image was also supplied. The resulting transformations were then concatenated with CIVET computed transformations between the T1w image and MNI space in order to form a single transformation from MNI space to DWI space. This single transform was used to warp cortical mid-surfaces from MNI space to DWI space. The transformed surfaces were then used to obtain mid-surface estimates of each of MD, FA, and RD, by supplying the transformed surfaces and each respective image to the volume_object_evaluate function (minc-toolkit v1.9.17). This process aims to measure DWI values at voxels which intersect with the mid-surface vertices, thus discarding values in white matter as well as grey matter superficial and deep to the mid-surface. Like CT and SA data, left and right midline data was masked out resulting in a total of 77122 vertex-wise data points for each of MD, FA, and RD.

### 2.4 Identifying Components using Non-negative Matrix Factorization

We used non-negative matrix factorization (NMF) to identify multimodal structural components, motivated by previous work from our group deriving a multi-modal parcellation of the human hippocampus (R. Patel et al., 2020). NMF is a matrix decomposition technique which decomposes an input matrix into two matrices containing components and weights, respectively. In the context of this analysis, NMF identifies regions of the brain where inter-individual macro- and microstructural variation is observed (spatial components) as well as each individual’s macro- and microstructural profile in a given component (subject weightings). Together, these outputs localize individual variability to specific brain regions in a data-driven manner. As the name suggests, NMF requires non-negativity in both inputs and outputs, leading to an additive parts-based representation (D. D. Lee & Seung, 1999). Given an input matrix of dimensions m x n, NMF outputs a component matrix W (m x k), and a weight matrix H (k x n). The number of components, k, is defined by the user. Each component describes a distinct multimodal covariance pattern across the input imaging data, with the W component matrix identifying the spatial location of the component (vertices with higher component scores represent where the component is located), and the H weight matrix quantifying the interindividual variability within the identified spatial region for each of the 5 input MRI metrics.

In this implementation, the NMF input matrix is constructed by stacking the vertex x subject matrices of each microstructural metric together. For each of the 5 metrics, vertex-wise data from all subjects is concatenated to form 5 separate matrices of dimensions 77122 rows x 398 columns (77122 vertices, 398 subjects). At each vertex and for each metric, we model out the effect of scanner using linear regression. Prior to concatenating individual metric matrices together, a per-vertex z-scoring is applied to each metric to account for differences in magnitude. This decision removes regional differences in structure, but instead emphasizes and focuses our analysis on interindividual differences.The resulting 5 matrices are then stacked side by side to form a matrix containing z-scored and residualized data with dimensions 77122 rows and 1990 columns (77122 vertices, 398 subjects * 5 metrics = 1990 columns). This matrix is shifted by its minimum value to create a non-negative input matrix for NMF. We used sklearn (version 0.23.1) to implement NMF with a non-negative singular value decomposition initialization to improve sparsity (Boutsidis & Gallopoulos, 2008). Number of components was selected through a split half stability analysis and a balance of spatial stability and model reconstruction accuracy (R. Patel et al.,2020).

The use of NMF in this study creates a two-step analysis procedure - MRI data is first input into NMF with the resulting components serving as input to the PLS analysis. It’s justification is three-fold. First, NMF provides an efficient form of dimensionality reduction of the input brain data. After extracting vertex-wise values of each MRI metrics, a total of 385610 measurements exist per subject (77122 vertices x 5 metrics). In light of potential concerns regarding the use of multivariate brain-cognition techniques in which the number of features vast outnumbers the number of subjects (Helmer et al., 2021; Marek et al.,2020), use of NMF significantly reduces the number of features per subject to help guard against concerns of latent variable stability. Second, NMF provides a means of multimodal data fusion. By inputting all metrics into NMF, the covariance across all metrics is considered simultaneously, a key point for fully taking advantage of multimodal data and its complementary nature (Groves et al., 2012). Finally, NMF has shown the ability to act as a detector of biologically relevant data in a number of neuroimaging applications (Nassar et al., 2018; Sotiras et al., 2015, 2017; Varikuti et al., 2018). By choosing to normalize data on a per-vertex basis as described above, we take advantage of this variance detection ability by restricting NMF to identify covariance patterns purely related to interindividual variability, as opposed to also including intraindividual (i.e. regional) variability related to regional differences in cortical structure. We thus constrain the downstream PLS analysis to focus on brain-cognition relationships in which the “brain” data represents, as best as possible, purely interindividual variability of cortical structure.

### 2.5 Cognitive Function Trajectories

We used 5 tests to measure cognitive performance. These include semantic fluency (in one minute, recall as many animals as possible), lexical fluency (in one minute, recall as many words starting with “S” as possible), short term memory (20 word free recall, recall within two minutes), inductive reasoning through the Alice-Heim 4-I (AH4) test (Heim, 1970), and vocabulary using the Mill Hill test (Raven,1958; Singh-Manoux et al., 2012). We included the total AH4 score (inductive reasoning) as well as mathematical and verbal reasoning sub scores, giving a total of 7 cognitive scores. For each score, a linear mixed effects model was performed with an interaction of baseline age and time since baseline as a fixed effect, a random slope of time since baseline, and random intercept for each subject (1).

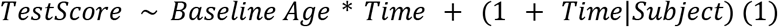

Models were implemented in R (version 3.6.3) using the nlme (version 3.1-149) package and implemented continuous autoregressive moving-average correlation structure to consider correlations between repeated measures on the same individual. Importantly, cognitive test data prior to the MRI time point (Wave 5-11 but excluding Wave 12) was included in the linear mixed effects modelling. For each model we extracted subject-specific intercepts, as well as the slope (i.e. coefficient) of the time variable (using the R coef() function). These values are a summation of the overall fixed effects and subject-specific random effects (De Jager et al., 2012), thus representing individual differences in modelled baseline performance (intercept) and impact of time (slope).

### 2.6 Partial Least Squares

To investigate structural-cognition relationships, we performed a brain-cognition partial least squares analysis (PLS). PLS is a multivariate technique which aims to maximize the covariance between two sets of variables (Krishnan et al., 2011; McIntosh & Lobaugh, 2004; McIntosh & Mišić, 2013). In this implementation, brain variables correspond to a 398 x 50 matrix containing NMF weightings of each subject within each of 10 components, for each of 5 MRI metrics. Cognitive data corresponds to a 398 × 14 matrix containing intercept and slope measures for each subject, for each of the 7 cognitive test scores. PLS outputs orthogonal latent variables (LV), each describing an independent pattern of covariance between NMF weights and cognitive intercepts and slopes. Each brain-cognition LV includes a singular value used to measure the proportion of total covariance explained. Statistical significance of an LV is assessed using permutation testing (n=10000), which develops a null distribution of singular values from which a non parametric p-value of each singular value in the original, non permuted data is computed. Bootstrap resampling is used to generate distributions of the singular vector weights of each brain and cognition variable, which enables identification of a confidence interval associated with each brain and cognition variables contribution to a given LV. For brain variables, the bootstrap resampling ratio (BSR) is computed as the ratio of the singular vector weight from the original run over the standard error of the weight derived from the corresponding bootstrap distribution. A BSR of high magnitude (BSR can be positive or negative) thus describes a brain variable with a strong and consistent contribution to an LV. We used a threshold of +/- 1.96 as a cutoff, analogous to a 95% confidence interval, such that only brain variables with a BSR magnitude above 1.96 are interpreted as contributing to an LV. (Krishnan et al.,2011; McIntosh & Lobaugh, 2004; Nordin et al., 2018; Zeighami et al., 2017). Finally, we obtained “brain” and “cognition” scores for each individual via multiplication of the output saliences and the original subject data. This outputs a brain score as well as a cognition score for each individual, and for each LV, which describes the degree to which a given individual expresses the pattern of covariance described by an LV (Zeighami et al., 2017). Matlab R2016a was used to perform the analysis along with the PLS package created by the Rotman Research Institute (http://pls.rotman-baycrest.on.ca/source).

### 2.7 Predicting Future Cognition Using Brain-Cognition Latent Variables

To further investigate the biological significance of the LVs, we performed linear models to examine the relationship between LV cognition scores and future cognitive performance (i.e. cognitive scores at Wave 12). For each cognitive score, we modelled Wave 12 performance as a function of LV1 and LV2 cognition scores while covarying for age, sex and years of education using a model of the form

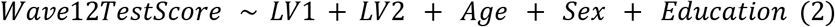

Where LV1 and LV2 terms represent cognition scores of each individual in a given LV. We performed the above model for 6 of the 7 test scores used in the PLS analysis described in section 2.6 (Mill Hill test was not conducted at Wave 12 and thus is not included in this analysis). Given the count nature and skew of cognitive test scores, a square root transformation followed by inverse normal transformation was applied to these data prior to linear modelling. Terms of interest included the LV1, LV2 terms and their interaction, assessed at a bonferroni threshold of 0.0083 (0.05 / 6 tests). To complete our characterization of LV cognition scores, we repeated this analysis using untransformed test score data and assessed relationships between LV cognition scores and each of age, MOCA status, sex, and education at the Imaging Wave.

### 2.8 Data and Code Availability

The study follows Medical Research Council data sharing policies (https://mrc.ukri.org/research/policies-and-guidance-for-researchers/data-sharing/). In accordance with these guidelines, data from the Whitehall II Study and the Imaging Sub-study are accessible via a formal application on the Dementias Platform UK portal (https://portal.dementiasplatform.uk/). Code used in this analysis is available at https://github.com/raihaan/micro-cog-nmf.

## 3.0 Results

### 3.1 Sample

The final analysis sample included 398 individuals who passed quality control for motion and cortical thickness processing, and had whole brain DWI available (mean age = 69.5 years ± 4.2, 92 females (23%), mean education years = 14.2 ± 3). Comparison of the analysis and initial samples is shown in Table 1. For further details on sample selection, see SI Methods and Figure S1.

**Table 1:** Demographic characteristics for the analysis and starting samples including mean and standard deviations for age, years of education, and MOCA score at the MRI Wave. Number of women, as well as number of individuals with MOCA score >=26, indicative of no major cognitive impairments, is shown. Statistical tests (t-test or chi-squared) show no differences between the analysis sample and full starting sample.

|  | **Analysis sample**<br>**(N=398)** | **Starting sample**<br>**(N=775)** | **Test Statistic** | ***p*** |
| --- | --- | --- | --- | --- |
| Age (years) | 69.52± 4.9 | 69.81±5.19 | $t = 0.95$ | 0.35 |
| Education (years) | 15.84±3.54 | 15.72±3.53 | $t = -0.58$ | 0.57 |
| MOCA score | 27.29±2.23 | 27.18±2.26 | $t = -0.80$ | 0.42 |
| Women, N (%) | 92 (23.1%) | 150 (19.3%) | chi-squared<br>= 2.27 | 0.13 |
| MOCA≥26, N (%) | 322 (80.9%) | 614 (79.2%) | chi-squared<br>= 0.46 | 0.49 |
| N (%) of participants scanned on Verio MRI scanner | 281 (70.6%) | 552 (71.2%) | chi-squared<br>= 0.49 | 0.82 |

### 3.2 Non-negative matrix factorization identifies 10 stable structural components

NMF analysis was performed on an input matrix consisting of cortical measurements of all subjects CT, SA, MD, FA, and RD data (R. Patel et al., 2020; Sotiras et al., 2015). Split half stability analysis (R. Patel et al., 2020) identified 10 components as a suitable balance between spatial stability and reconstruction error (see SI Methods and Figure S2). The 10 spatial components and associated weightings are displayed in Fig 2A and 2B, respectively. Each component identifies a group of vertices which share a covariance pattern for CT, SA, MD, FA, and RD. The components are largely bilateral and non-overlapping, and their regional descriptions and naming conventions are described below.

1. Component 1: (Fronto-Temporal) is localized in the superior frontal and posterior temporal regions.
2. Component 2: (Motor) is localized to primary and supplementary motor cortices, with some spread to adjacent posterior frontal and superior parietal regions.
3. Component 3: (Visual) is strongly localized in the medial and lateral occipital lobe, as well as the cingulate cortex and inferior temporal lobe.
4. Component 4: (Parietal) occupies most of the parietal cortex, with some spread to the lateral temporal regions.
5. Component 5: (Inferior Frontal) is most prominent in the inferior, medial frontal lobe, but also shows some presence in the inferior temporal lobe, anterior cingulate regions, and inferior lateral frontal lobe.
6. Component 6: (Anterior Frontal) occupies the anterior frontal regions as well as the temporal pole.
7. Component 7: (Cingulate) occupies much of the midline regions but with a strong preference to the cingulate cortex and shows some spread to insular cortices.
8. Component 8: (Postcentral) is heavily localized to the postcentral gyrus but shows considerable presence in the lateral inferior frontal lobe and superior temporal gyrus.
9. Component 9: (Right lateralized) is the only component showing a laterality effect, including bilateral medial parietal anterior temporal regions, but most prominent in right superior temporal and lateral inferior frontal regions.
10. Component 10: (Temporal Pole) is most prominent in the temporal pole, but also shows some presence in medial temporal and ventromedial frontal areas.

**Figure 2.**
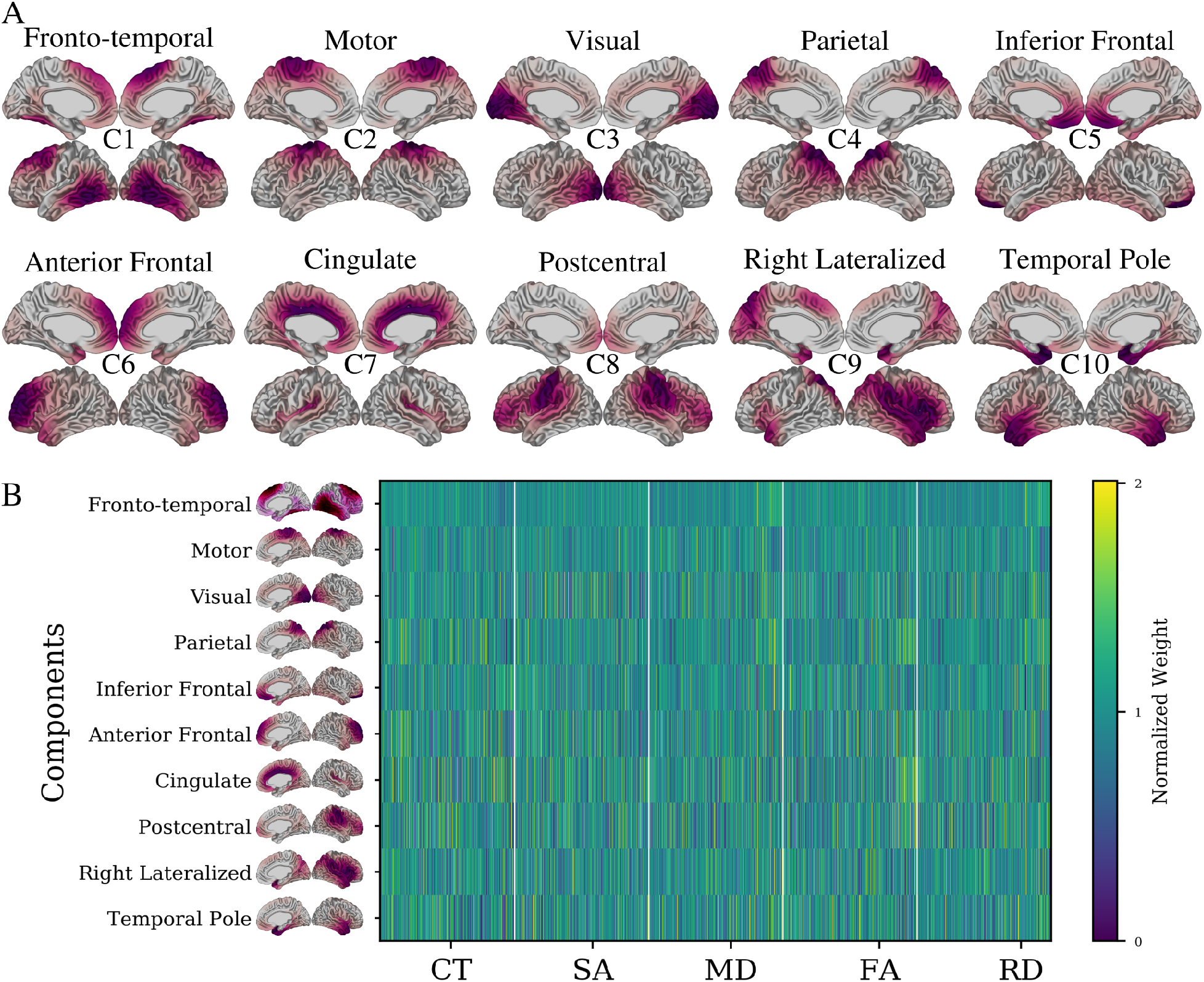
10 structural components derived from the NMF decomposition. A) Cortical mappings of each of the spatial patterns for each of the 10 components. For each component, lateral and medial views of both left and right hemispheres are shown. Components are identified using the putative descriptors from the main text (e.g., Fronto-temporal) as well as lettering at the centre of each set of surface views (e.g., C1). Red areas indicate vertices loading heavily onto a particular component (thresholded at 25% to max value). Each component identifies a selection of vertices sharing a structural variance pattern. Together, components cover the entire brain, are largely bilateral (with exception of component 9) and are not spatially overlapping. B) Subject-specific weightings associated with each of the displayed 10 components. Each row corresponds to a specific component’s NMF weightings for each subject-metric combination, describing the CT, SA, MD,FA and RD patterns of each subject in each component. Together these two outputs describe the morphological patterns of each subject within each spatial component. Each element of the matrix is displayed as a fraction of its row mean, such that values below 0 indicate a below average weight for a given component and vice versa.

Each of the 10 components represents a set of vertices sharing a covariance pattern across the input imaging data. Recalling the per-vertex normalization procedure described in Section 2.4, which prioritized interindividual variability, these components represent between-subjects variability derived from multimodal MRI measures of cortical structure. Each component can be described via a spatial pattern (Figure 2A) as well as a set of weightings for each subject and MRI metric (Figure 2B) quantifying the variability across individuals. To further probe the significance of these patterns in the context of cognitive decline, these subject weightings were used as input to PLS analysis.

### 3.3 Specific Patterns of Cortical Morphology Relate to Baseline and Longitudinal Cognitive Function

We next related the variation in cortical structure captured by the NMF subject weightings to variability in cognitive performance over time. For each participant we derived the intercept and slope for the change in performance across each of seven cognitive tests: lexical and semantic fluency, short-term memory, verbal, mathematical and inductive reasoning, and vocabulary (SI Results Table S1). The intercept describes the estimated baseline (i.e., mid-life) performance while the slope describes the linear rate of change in performance over time (i.e., from mid-life to late-life). We performed a brain-cognition PLS with NMF component weightings as “brain” data and intercept and slope measurements as “cognition” data. PLS analysis identified two significant latent variables (LVs), explaining 56.9% and 19.7% of shared brain-cognitive covariance respectively. Each LV identifies a distinct pattern of longitudinal cognitive trajectories across the 20-year follow up that relate to patterns of late-life structural characteristics (Figure 3). As the LVs contain a mix of cross sectional and longitudinal measures, in the descriptions below the use of the words “decline”, “increase”, and “decrease” specify changes over time. Conversely, when describing the brain features of each LV (derived from cross sectional MRI), use of the words “higher” and “lower” is instead used and pertains to relatively higher or lower MRI measures in relation to the comparison group at study.

**Figure 3:**
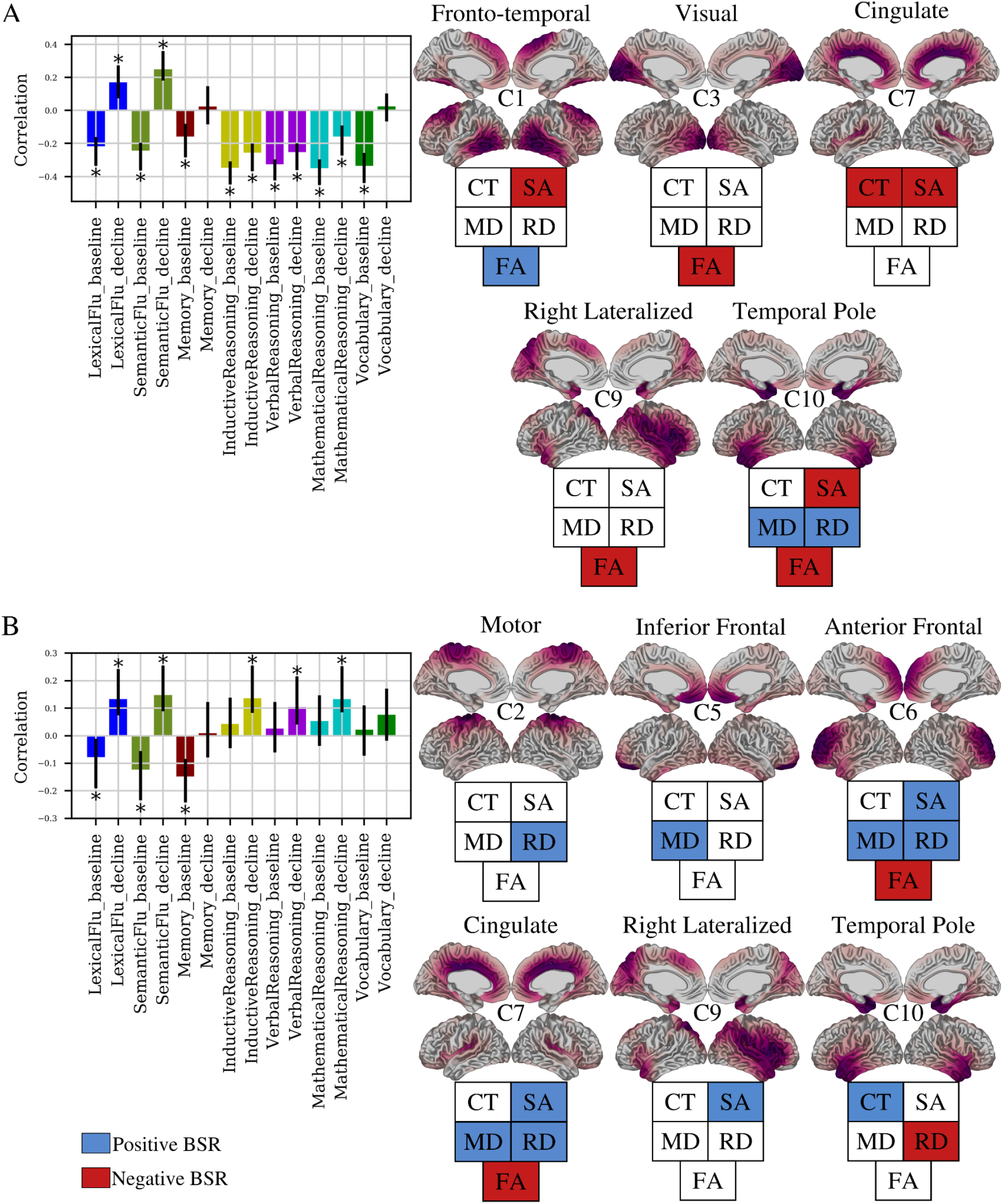
Two brain-cognition latent variables identified by PLS. PLS identified two (LV1: A, LV2: B) significant latent variables (p<0.05), each identifying a pattern of covariance between NMF weights and cognitive intercept and slopes. Bar plots describe the contribution of cognitive variables. The y-axis denotes correlation of each cognitive variable in a LV. Error bars denote 95% confidence interval, only variables with non-zero confidence interval are described as contributing to a LV (marked with *). For each bar plot, cortical maps (right) show spatial patterns of the components contributing to the LV. The fingerprint of each map describes whether a given metric is positively (blue) or negatively (red) associated to the cognitive pattern shown in bar plots.

To interpret the brain-cognition patterns identified by PLS, we have presented a cognitive plot and associated brain maps. The cognitive plot shows the correlation of each brain variable with a given LV. Two variables with the same direction (e.g. both positive) are identified to covary together, while those with opposing directions (e.g. positive and negative) covary negatively. If the error bars of a variable cross zero, the contribution of this variable to the LV is not deemed to be reliable and is excluded. For example, in LV1, baseline performance on fluency (negative correlation) covary negatively with fluency decline measures (positive correlation) but covary positively with memory baseline performance. To identify the brain variables involved with the cognitive pattern described by the LV, the BSR of each brain variable can be used. Specifically, those variables with a positive BSR above the BSR threshold are positively associated with the cognitive pattern, while those with negative BSR below the BSR threshold are negatively associated. For example, the cognitive features of LV1 are positively associated with FA in component 1 (blue, positive BSR), and negatively associated with SA in component 1 (red, negative BSR).

LV1 (p = 0.0042) describes a pattern in which low baseline performance across all tests, accelerated 20-year decline in inductive, verbal, and mathematical reasoning abilities, but slow 20-year decline in verbal and semantic fluency are associated with low late-life SA across the brain, lower CT (cingulate/insular), lower FA (visual, temporal, right lateral), higher MD and RD (temporal pole), and higher FA in posterior temporal and superior frontal regions. (Figure 3A, Table 2). LV2 (p = 0.0003) describes a pattern in which low baseline performance in each of lexical fluency, semantic fluency, and short term memory, but slower decline in each of lexical fluency, semantic fluency, inductive reasoning, verbal reasoning, and numeric reasoning is associated with high SA across the brain, higher CT and lower RD in the temporal pole, higher MD in inferior frontal, anterior frontal, and cingulate cortices, higher RD in motor, anterior frontal, and cingulate cortices, and lower FA (anterior frontal, cingulate/insular) (Figure 3B, Table 2).

**Table 2:**
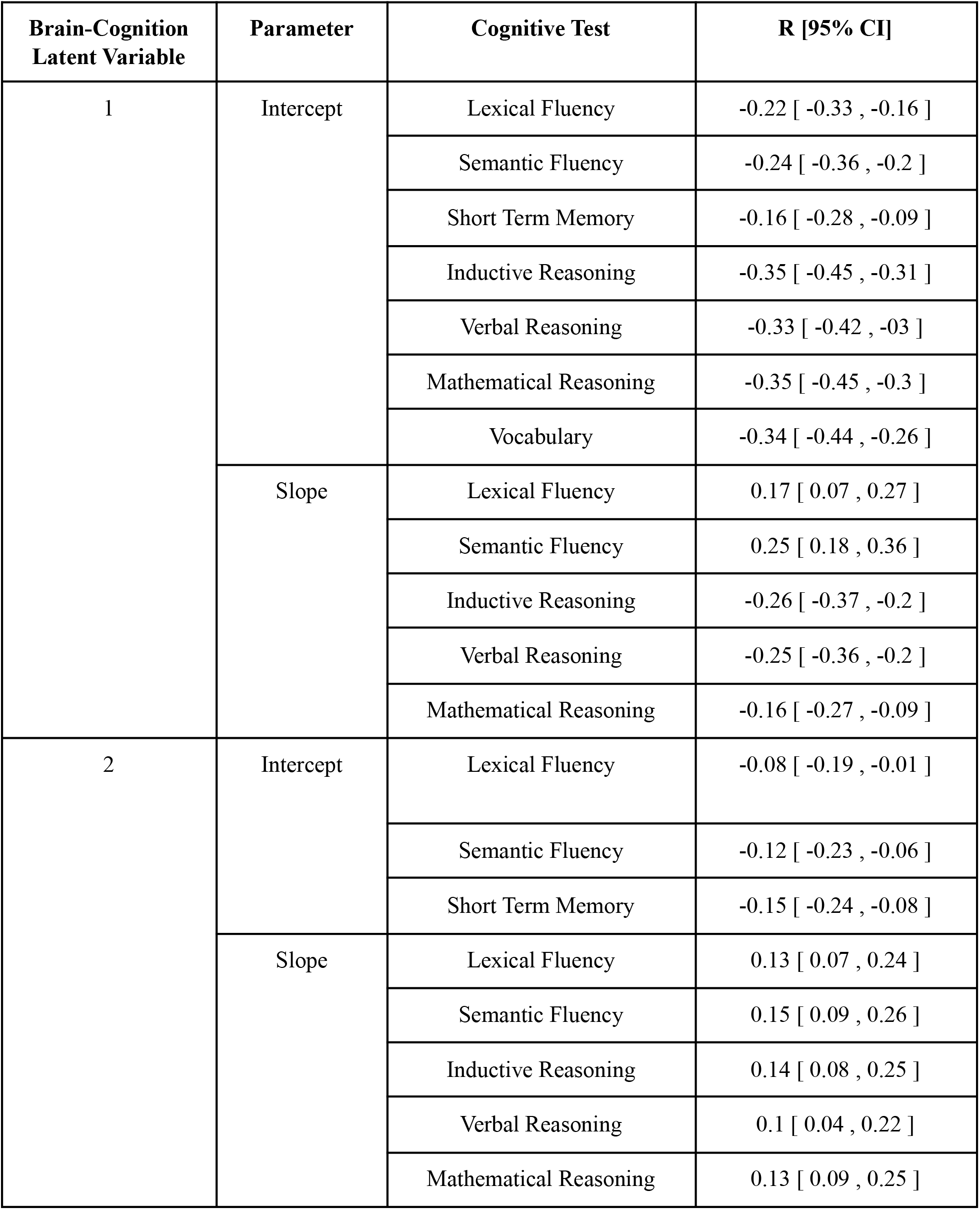
Correlations of the contributing cognitive variables to each latent variable

| Brain-Cognition<br>Latent Variable | Parameter | Cognitive Test | R [95% CI] |
| --- | --- | --- | --- |
| 1 | Intercept | Lexical Fluency | -0.22 [ -0.33 , -0.16 ] |
|  |  | Semantic Fluency | -0.24 [ -0.36 , -0.2 ] |
|  |  | Short Term Memory | -0.16 [ -0.28 , -0.09 ] |
|  |  | Inductive Reasoning | -0.35 [ -0.45 , -0.31 ] |
|  |  | Verbal Reasoning | -0.33 [ -0.42 , -0.3 ] |
|  |  | Mathematical Reasoning | -0.35 [ -0.45 , -0.3 ] |
|  |  | Vocabulary | -0.34 [ -0.44 , -0.26 ] |
|  | Slope | Lexical Fluency | 0.17 [ 0.07 , 0.27 ] |
|  |  | Semantic Fluency | 0.25 [ 0.18 , 0.36 ] |
|  |  | Inductive Reasoning | -0.26 [ -0.37 , -0.2 ] |
|  |  | Verbal Reasoning | -0.25 [ -0.36 , -0.2 ] |
|  |  | Mathematical Reasoning | -0.16 [ -0.27 , -0.09 ] |
| 2 | Intercept | Lexical Fluency | -0.08 [ -0.19 , -0.01 ] |
|  |  | Semantic Fluency | -0.12 [ -0.23 , -0.06 ] |
|  |  | Short Term Memory | -0.15 [ -0.24 , -0.08 ] |
|  | Slope | Lexical Fluency | 0.13 [ 0.07 , 0.24 ] |
|  |  | Semantic Fluency | 0.15 [ 0.09 , 0.26 ] |
|  |  | Inductive Reasoning | 0.14 [ 0.08 , 0.25 ] |
|  |  | Verbal Reasoning | 0.1 [ 0.04 , 0.22 ] |
|  |  | Mathematical Reasoning | 0.13 [ 0.09 , 0.25 ] |

In summary, each brain-cognition LV described mixed patterns of cognitive decline and maintenance, as well as both positive and negative associations with cortical structure indices. This suggests that rather than cognitive decline being uniformly negatively associated with cortical structure, individuals display degrees of cognitive decline and maintenance across diverse cognitive functions that are both positively and negatively associated with cortical structure.

### 3.4 Brain-Cognition Latent Variables Predict Cognitive Performance at Future Timepoint

To investigate the biological significance of the brain-cognition relationships identified by the two distinct LVs, we assessed how expression of LVs predicted cognitive performance at a future time point (Wave 12). In doing so, we were particularly interested in seeing if baseline performance may be better predictors of future cognition than measurements of decline. Throughout the LV patterns, there are varying combinations of baseline and decline. For example, an individual with high LV1 expression performed poorly at the baseline measurement on tests of fluency but exhibited slower decline. A natural follow up to this observation is if this individual performs better in the future than their counterpart, who showed the inverse pattern of higher baseline performance but accelerated decline. Within each LV, a cognition score was computed for each subject by projecting PLS derived saliences on the input cognitive data. These scores describe the degree to which an individual expresses the cognitive phenotype described by LV1. For example, LV1 describes low semantic fluency baseline performance but slower decline measures (Figure 3A). Thus, an individual with a high LV1 cognition score would be expected to have low semantic fluency at baseline but a slower decline. This is illustrated by the plots in Figure 4A, which show a histogram of LV1 cognition scores and a strong negative association to semantic fluency baseline and positive association to semantic fluency decline (Figure 4A).

**Figure 4:**
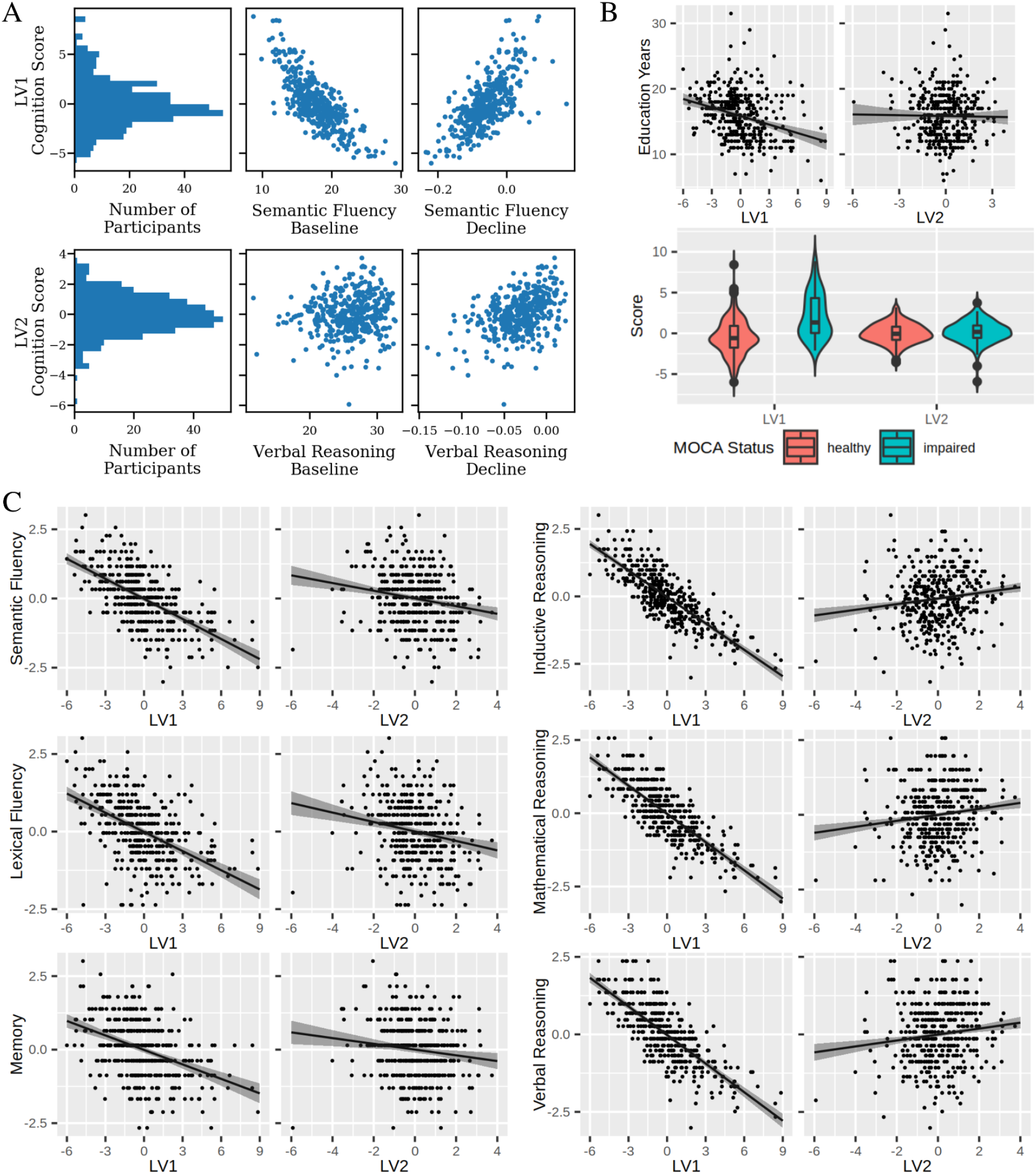
Individual expression of PLS derived brain-cognition latent variables is indicative of cognitive status and future performance. We used each subject’s expression of LV1 and LV2 to predict their cognitive performance at a future wave. A) An illustration of the utility of LV cognition scores. Top row: a histogram shows the distribution of LV1 cognition scores, which quantify the degree to which each subject expresses the phenotype described by LV1. Plots of LV1 versus semantic fluency baseline and decline measures show positive and negative relationships, respectively. These match that described in LV1 in Figure 3, demonstrating that LV cognition scores can be used to describe the cognitive phenotype of subjects. Bottom row: equivalent plots for LV2 including the histogram, and plots of LV2 cognition scores vs verbal reasoning baseline (no relation) and verbal reasoning decline (positive relation). B) LV1, but not LV2, cognition scores relate to key demographic variables. Linear modelling of years of education as a function of LV scores showed a negative relationship with LV1 (β = −0.43, p<0.01), but no relationship with LV2 (β = −0.04, p>0.05). Similarly, t-test of mean score between healthy and impaired MOCA groups showed significant differences for LV1 (t(98.5)=7.9, p<0.01) but not LV2 (t(101.9)=0.79, p>0.05). C) Effects plots of each Wave 12 test vs. LV1 and LV2, respectively. Effects plots, created using the effects() R function, plot the predicted test score at various levels of either LV1 or LV2 while assuming population averages for other covariates. Shaded areas represent confidence intervals around the estimate. Table 3 displays statistics from the linear models used to assess these relationships, each of which was significant at a bonferroni threshold of p < 0.0083 (0.05/6).

Higher expression of the LV1 phenotype was associated with having fewer years of education (β=-0.43, p<0.01) and lower MOCA scores (i.e. greater cognitive impairment) (t(98.5)=7.9, p<0.01, Figure 4B) at the Imaging Wave. However, LV2 scores were not related to either MOCA (t(101.9)=0.79, p>0.05) or education (β=-0.04, p>0.05) (Figure 4B). Both LV1 and LV2 showed sex differences, with females having higher LV1 but lower LV2 scores in comparison to males (SI Results and Figure S3). Neither LV1 or LV2 showed any relationship to age (SI Results and Figure S3).

We next performed linear models to examine the relationship between LV scores and Wave 12 cognitive performance. Table 3 lists standardized coefficients and p values of these analyses, assessing significance at Bonferroni corrected threshold of p<0.0083 (0.05/6). We observed that high expression of LV1 scores was associated with lower performance on each test. Conversely, higher expression of the LV2 was associated with lower Wave 12 performance of semantic fluency, lexical fluency, and memory tests but better performance on inductive, mathematical, and verbal reasoning tests. Thus, we find both LV1 and LV2 are predictive of future cognitive performance in unique and distinct ways. A second model was run with time interval between waves as an additional covariate which showed similar results (Supplementary Table S3).

**Table 3:** Statistical results from linear models run for Wave 12 performance on each of 6 cognitive tests. Each model had the form *Test* ~ *LV*1 + *LV*2 + *Age* + *Sex* + *Education*. Standardized beta coefficients and 95% confidence intervals are reported, where the standardized coefficient represents the change in wave 12 test scores for every one unit increase in LV scores, in units of standard deviation. Bold p-values are deemed significant at a Bonferroni-corrected threshold of p<0.0083.

| Test | Term | Standardized $\beta$ [95% CI] | p |
| --- | --- | --- | --- |
| Semantic Fluency | LV1 | -0.61 [-0.69 -0.53] | <0.001 |
|  | LV2 | -0.18 [-0.26 -0.11] | <0.001 |
| Lexical Fluency | LV1 | -0.52 [-0.61 -0.43] | <0.001 |
|  | LV2 | -0.20 [-0.28 -0.12] | <0.001 |
| Memory | LV1 | -0.42 [-0.51 -0.33] | <0.001 |
|  | LV2 | -0.13 [-0.22 -0.04] | 0.003 |
| Inductive Reasoning | LV1 | -0.82 [-0.88 -0.77] | <0.001 |
|  | LV2 | 0.13 [0.08 0.18] | <0.001 |
| Mathematical Reasoning | LV1 | -0.81 [-0.87 -0.76] | <0.001 |
|  | LV2 | 0.13 [0.08 0.19] | <0.001 |
| Verbal Reasoning | LV1 | -0.78 [-0.84 -0.72] | <0.001 |
|  | LV2 | 0.13 [0.07 0.18] | <0.001 |

## 4.0 Discussion

In this study, we used a data-driven approach to identify brain-cognition latent variables of covariance linking individual variation in cognitive decline over a period of 20-years from mid-to-late life, with later-life patterns of cortical structure. We present several novel results. First, we demonstrate that the association between cognitive decline and cortical structure as assessed by multivariate data-driven methodologies is complex and heterogenous. Related univariate studies have yielded positive brain-cognition associations in a relatively consistent fashion (Oschwald et al., 2019). Further, purely hypothesis driven approaches may be constrained by prior definitions and categorizations, limiting the detection of heterogeneity in the sample (Habes et al., 2020). By operating with a hypothesis free and data-driven approach, here we instead demonstrate brain-cognition relationships in which the directionality of brain-cognition, as well as cognition-cognition relationships is more variable. For example, LV2 shows that in our sample, a relative maintenance over time of reasoning abilities covaries with lower baseline performance in fluency and memory tests, and that these cognitive characteristics are associated with higher cortical area but also higher diffusivity in the anterior frontal regions. Second, the simultaneous analysis of multimodal MRI measures using NMF better enables us to uncover global and distributed networks in which cortical macro- and microstructural features are associated with cognition. A common approach thus far has been to relate a single measure of brain structure with one or more cognitive tests (Oschwald et al., 2019), identifying localized regions associated with cognition. Our approach instead investigated vertex-wise measurements of the entire cortex and identified networks of brain regions with shared covariance patterns (Sotiras et al., 2015). We include a range of macro- and microstructural level metrics, increasing the sensitivity of our analysis to complementary underlying neurobiological mechanisms (Tardif et al., 2016). The resulting analysis enabled us to identify large-scale distributed brain regions which covary with cognitive maintenance and decline, suggesting that single region approaches may be obscuring the importance of numerous other brain regions. Third, we show that individual variation within these LVs is predictive of future cognitive performance, helping to understand the biological significance of these distinct brain-cognition LVs. From a technical perspective, our study adds to a growing body of evidence showing that multivariate data-driven analyses offer the ability to uncover distributed networks of brain regions linked to cognition (Assem et al., 2020; Bassett et al., 2020;Bassett & Sporns, 2017; Pichet Binette et al., 2020; van den Heuvel & Sporns, 2013). From a biological perspective, our study emphasizes the importance of mid-life cognitive health when considering late-life cognitive performance.

### 4.1 Brain-Cognition latent variables include a mixture of maintenance and decline

The two brain-cognition relationships identified in this work contain a mixture of positive and negative features across both brain and cognition. The first brain-cognition LV describes a pattern of low midlife performance across all tests, accelerated decline in reasoning, but relatively maintained fluency associated with a multimodal pattern of low SA in all areas of the brain except for the primary and supplementary motor cortices, low CT in cingulate and insular cortex, high diffusivity in the temporal pole, low FA in visual, temporal, and right lateral cortex but high FA in superior frontal and lateral temporal regions. Meanwhile, the second brain-cognition LV links low baseline fluency and memory performance, but slower fluency and reasoning declines with higher diffusivities in nearly all regions except occipital cortex, high surface area in cingulate, insular and right lateral areas, high temporal pole thickness, and low FA in frontal, cingulate and insular regions. Thus, across both brain and cognition, we observe a mix of what studies have traditionally characterised as adaptive and maladaptive characteristics. Previous findings have linked performance in fluency tests to fronto-temporal regions (Rodríguez-Aranda et al.,2016; Zhang et al., 2013), and potentially parietal cortex as well (Rodríguez-Aranda et al., 2016).

Similarly, frontal cortex has been highlighted as playing a key role in reasoning abilities (Mole et al.,2021). These findings are in line with general theories attributing cognitive decline to impairments of processing speed and executive function in which the frontal lobes play a prominent role (Jagust, 2013). Our findings instead suggest a more distributed and variable pattern, in which not only other regions such as the visual (LV1) and motor cortex (LV2) are involved, but the directionality of associations may not be as straightforward as simple positive associations. While it may be tempting to infer patterns of overall cognitive or neuroanatomical maintenance, our work suggests that by refraining from a priori definitions of maintenance or decline groups, including baseline and decline measures across a range of tests, and analysing multimodal indices of cortical structure simultaneously, we can identify subtle, complex brain-cognition relationships which show a mix of maintenance and decline features across both cognitive and anatomical measures. These findings thus discourage the use of a ‘one size fits all’ approach, and instead encourage the consideration of cognitive domains and regional, multivariate anatomy at the individual level.

While we cannot derive direct mechanistic inferences, our results also shed light on the neurobiological underpinnings of the two latent variables. Histological evidence links CT reductions in old age with decreased dendritic arborization (Esiri, 2007; Goriounova et al., 2018). A recent application of virtual histology supports this, having found cortical thinning is related to increased expression of CA1 pyramidal cell gene sets enriched with processes related to dendritic arborization approach correlated longitudinal CT changes with cell type expression levels (Y. Patel et al., 2021; Vidal-Pineiro et al., 2020). Corresponding studies of SA remain uninvestigated, though see our discussion in Section 4.2 for more on our SA findings. DWI metrics may relate to a range of age-related alterations such as decline in small diameter myelinated fibers, alterations in the myelin sheath, and inflammatory processes (Beaulieu, 2002;Marner et al., 2003). Increased diffusivity is a common finding in aging studies, and may relate to enlarged interstitial spaces or axon swelling (Benedetti et al., 2006; Chad et al., 2018; Madden et al.,2012). Rodent studies incorporating imaging and histology demonstrate that demyelination of axons leads to increased RD (Song et al., 2002, 2004; Sun et al., 2008; Tuor et al., 2014). Common interpretations of increased FA is related to the presence of a preferential fiber orientation, and of high myelin content (Alexander et al., 2007; Basser & Ozarslan, 2010; Beaulieu, 2002; Lazari & Lipp, 2021; Le Bihan et al.,2001; Mancini et al., 2020), however these interpretations are based on the study of FA in white matter where it is accepted that orientation of myelinated axons represents the dominant source of anisotropy (Edwards et al., 2018). In the cortex, interpretation of anisotropy is more challenging and less explored (Edwards et al., 2018; Lampinen et al., 2019). Though evidence of anisotropy in the cortex has been demonstrated (Aggarwal et al., 2015; McNab et al., 2013; Truong et al., 2014), FA is generally lower in cortex than in white matter (Assaf, 2019; McNab et al., 2013) and may be driven by a number of factors including unmyelinated axons (Assaf, 2019; Leuze et al., 2014; Nair et al., 2005), dendrites (Jespersen et al., 2007), and cell bodies (Edwards et al., 2018). Thus we restrict our interpretation of cortical FA changes to decreased FA potentially reflecting general degeneration as well as lower myelin content. We hypothesize that LV1 has a neurobiological pattern of widespread reductions in cortical area, dendritic branching, demyelination, and axonal degeneration. A surprising feature of LV1 is the higher FA in the lateral and temporal regions of component 1. Higher FA in aging populations has been hypothesized to represent a degeneration of secondary fiber populations rather than increases in fiber coherence (Douaud et al., 2011; Miller et al., 2016). We consider this a tentative explanation of our FA findings in LV1, though based on the complexities of FA interpretation in cortical grey matter discussed above we avoid strong interpretations of differential fiber orientation based on our findings. Meanwhile, LV2 is associated with near global demyelination and axonal degeneration, but relatively maintained cortical area in cingulate, insular, right lateral cortex and dendritic morphology in the temporal pole. However, while plausible, these interpretations are complicated by the fact that each MRI metric is sensitive to a large range of cellular level alterations (Tardif et al., 2016; Zatorre et al., 2012) as well as the interrelated nature of the DWI metrics analysed (Madden et al., 2012). While joint analyses of all metrics, as is a focus of this work, may help alleviate some concerns (Assaf & Pasternak, 2008), caution is still warranted in absence of direct histological evidence.

### 4.2 Late Life Cognitive Performance is Driven by Mid Life Phenotypes

In our study sample and within the timeframe examined, the strongest predictor of later life (>65 years) cognitive performance across a range of tests was performance in midlife (at mean age of 40 years). We assessed the relationship between LV cognition scores and cognitive performance at a future time point and found that across all tests, individuals scoring high on LV1 performed worse than those who expressed the inverse cognitive pattern. In certain cases this is a straightforward result, as the cognition features of LV1 included either decreased baseline performance (memory) or both decreased baseline performance and accelerated decline (inductive, mathematical, and verbal reasoning). However, for tests of lexical and semantic fluency LV1 describes low baseline performance and slower decline. In this case, the inverse relationship between LV1 and future performance shows the relative maintenance of these abilities is not enough to compensate for lower initial performance. Individuals scoring high on LV2 performed worse on tests of fluency and memory, but better on tests of inductive, mathematical, and verbal reasoning compared to those with lower LV2 scores. Similar to LV1, LV2 describes low baseline performance and slower decline on both fluency measures, and the inverse relationship between LV2 and future performance shows maintenance is not enough to compensate for lower initial performance. For tests of inductive, verbal and mathematical reasoning, LV2 describes a relative maintenance without any baseline effects, in line with the positive association between LV2 and future performance on these tests.

Inclusion of both LV1 and LV2 scores in our models suggests that each of the brain-cognition LVs identified are distinct in their ability to predict future cognitive performance. That we covaried for impacts of age, sex, and education further supports the unique impact of each LV, over and above other factors considered. For tests of inductive, mathematical, and verbal reasoning, LV1 and LV2 describe opposing characteristics with LV1 showing low baseline performance and accelerated decline and LV2 showing a relative maintenance over time. When predicting future performance, we found that LV1 beta coefficients were consistently larger in magnitude compared to LV2, suggesting the low midlife characteristics of LV1 have a larger impact on future performance than the maintenance characteristics of LV2. This supports the notion that low midlife performance is not adequately countered by the identified longitudinal maintenance. This finding has relevance to the concept of cognitive reserve. High levels of cognitive reserve, often probed through proxies such as education or occupational attainment, have been strongly linked to better cognitive function in aging (Ewers, 2020). Whether this is driven by a maintenance of previously developed advantages, moderation of the effects of aging on cognition and hence cognitive decline, or a combination however remains unclear (Ewers, 2020; Soldan et al., 2020). Our findings altogether suggest that midlife differences in cognitive performance were the most prominent predictors. However, it may be that individuals in our study sample have not yet reached a point of drastically accelerated decline and therefore showed relatively less profound differences in cognitive trajectories (Gregory et al., 2017).

While the MRI data is cross sectional and collected only in late life, in the context of mid life dominance we find the widespread involvement of SA to be of particular interest. SA is commonly assessed in parallel with CT, though the contribution of each to cortical volume, as well as their neurobiological and genetic underpinnings (Panizzon et al., 2009; Rakic, 1988, 2000; Storsve et al., 2014), varies. For example, while each of SA, CT, and cortical volume decrease with age, CT and volume change are positively correlated while changes between CT and SA tend to be inversely related (Storsve et al., 2014). SA decreases during aging are also of smaller magnitude than those observed for CT (Storsve et al.,2014), and the primary determinant of SA, cortical column generation, occurs during prenatal and perinatal periods (Rakic, 1988, 2000). These findings suggest SA may be more temporally fixed than CT between mid and late life, and throughout the full lifespan. In this context, the pronounced influence of LV1 mid life cognitive performance and late life SA on late life cognition lends credence to a lifespan perspective in which developmental and mid-life events play a significant role in cognitive health in late life. In a unique study involving the Lothian Birth Cohort, positive cognitive ageing between childhood (age 11) and late life (age 73) was associated with higher SA in late life (Cox et al., 2018). In another unique study involving multiple samples, Walhovd et al. identified a large region of the cortex in which increased general cognitive ability was associated with increased surface area in a developmental sample (aged 4-12) and noted that this association persisted throughout the lifespan (Walhovd et al., 2016). While the above discussion focussed on SA, we also note the widespread involvement of FA. As described below, the involvement of SA is of particular interest due to its reported life course characteristics as well as relevant works assessing SA and cognition. At this point, the life course characteristics of FA and cognition are not as concrete and so we hesitate to expand further on FA involvement in this context. Our findings, along with others, support the idea that stable advantages may give certain individuals a ‘head start’ in terms of cognitive function in aging. They also support the need for early and mid-life preventative measures of cognitive decline to maintain cognitive performance in older ages, though the potential role of reserve mechanisms on cognitive decline warrant further investigation. The mixed, complex relationships identified also support the development of individualized preventative strategies. Interestingly, within a subset of the Whitehall II cohort, previous work has identified similar mid to late life relationships across different biological domains such that midlife cardiovascular health was predictive of indicative of cerebral hypoperfusion in late life (Suri et al., 2019).

### 4.3 Strengths and Limitations

A key strength of this study is the use of multimodal MRI data to characterize cortical morphology. Use of multiple MRI metrics provides complementary information regarding anatomical properties of the brain in comparison to unimodal analyses. We take this further by employing NMF to analyse multimodal data simultaneously to capture shared patterns of covariance across measures. This approach allows us to identify 10 major components which are spatially contiguous and highlight relevant regions in which cortical morphology varies across subjects. We relate this variability to longitudinal cognitive performance using PLS. This approach does not limit us to broad categorizations of decliners or maintainers, rather, we obtain continuous measures for each individual identifying the degree of expression of each of the identified brain-cognition latent variables. The use of multimodal data and longitudinal cognitive measures is made possible by the unique dataset analysed. However, our study is limited by a smaller sample size in comparison to large scale neuroimaging analyses. It may be that more brain-cognition relationships would be identifiable in a larger sample. Our study also lacks an out of sample validation, as the unique longitudinal data makes a compatible out of sample dataset difficult to find. Further, because of our sample size we elected not to hold out a portion of the dataset to perform an internal validation, and instead used the full dataset to improve the descriptive nature of our analysis at the cost of its predictive utility. Future work with independent validations are needed in larger longitudinal MRI datasets as they emerge. In addition to this, the Whitehall II Imaging Sub-study cohort contains a higher proportion of men compared to the general population and is relatively more educated. Thus, generalizability of our findings to the wider population is limited. While we were able to quantify 20-year cognitive trajectories, MRI data is currently available at a single time-point which limits investigations of longitudinal brain-cognition relationships. Furthermore, while a combination of structural and diffusion MRI was used to provide a more comprehensive assessment of cortical structure, the limitations of MRI, in particular its resolution in comparison to the neural substrates under study, preclude us from inferring the cellular mechanisms which may be at play (Tardif et al., 2016). The use of NMF in this study enables a multimodal fusion and dimensionality reduction simultaneously, though unlike similar techniques such as Principal Component Analysis, exact quantification of variance explained is unavailable. Finally, we focus this analysis on the neocortical grey matter which captures a distributed set of brain regions, but this decision makes our study blind to the role of the hippocampus and subcortical structures, both of which have been shown to be heavily involved in brain aging (Bartsch & Wulff, 2015; Bussy, Patel, et al., 2021; Bussy, Plitman, et al., 2021; Fjell, McEvoy, et al., 2014; Tamnes et al., 2013; Tullo et al., 2019; Walhovd et al., 2011). Re-examining this research question using macro- and micro-structural measures of these structures (similar to our previous work in healthy young adults (R.Patel et al., 2020; Robert et al., 2022), could provide a pathway forward for reconciling the role of these structures.

## 5.0 Conclusion

This work provides novel information on brain-cognition relationships in a healthy elderly population. We uncover complex brain-cognition relationships using an unbiased data-driven approach, free of a priori definitions of cognitive maintainers or decliners and including a rich and comprehensive longitudinal cognitive data and multimodal MRI measures. This supports future works including multimodal data as well as cognitive trajectories to capture the full range of brain-cognition relationships. We also find the largest determinant of late life cognition is mid life cognition, as opposed to the rate of decline over time. This, and the associated link with widespread surface area measurements, support early and mid-life preventative measures of cognitive decline.

## Acknowledgements

We would like to thank all contributors to the Whitehall II study. We thank all the participating civil service departments; the British Occupational Health and Safety Agency; the British Council of Civil Service Unions; all participating civil servants in the Whitehall II Study; and all members of the Whitehall II Study team at University College London who so helpfully collaborated with us. The Whitehall II Study team comprises research scientists, statisticians, study coordinators, nurses, data managers, administrative assistants, and data entry staff, who make the study possible. We would like to thank staff at the Wellcome Centre for Integrative Neuroimaging in Oxford who acquired scans, in particular Dr Nicola Filippini (DPhil) and research radiographers Michael Sanders, MSc, Jon Campbell, MMRTech, BcAppSc, Caroline Young, DCR(R), David Parker, BSc(Hons). Martin R. Turner, MA, MBBS, PhD, FRCP (Wellcome Centre for Integrative Neuroimaging, Oxford, United Kingdom), and his colleagues advised on incidental findings and taking over clinical responsibility for such participants.

Computations were performed on the Niagara supercomputer at the SciNet HPC Consortium (Loken et al., 2010; Ponce et al., 2019). SciNet is funded by: the Canada Foundation for Innovation; the Government of Ontario; Ontario Research Fund - Research Excellence; and the University of Toronto.

## References

Ad-Dab’bagh, Y., Lyttelton, O., Muehlboeck, J. S., Lepage, C., Einarson, D., Mok, K., Ivanov, O., Vincent, R. D., Lerch, J., Fombonne, E., & Others. (2006). The CIVET image-processing environment: a fully automated comprehensive pipeline for anatomical neuroimaging research. Proceedings of the 12th Annual Meeting of the Organization for Human Brain Mapping, 2266.

Aggarwal, M., Nauen, D. W., Troncoso, J. C., & Mori, S. (2015). Probing region-specific microstructure of human cortical areas using high angular and spatial resolution diffusion MRI. NeuroImage, 105, 198–207.

Alexander, A. L., Lee, J. E., Lazar, M., & Field, A. S. (2007). Diffusion tensor imaging of the brain. Neurotherapeutics: The Journal of the American Society for Experimental NeuroTherapeutics, 4(3), 316–329.

Andersson, J. L. R., Skare, S., & Ashburner, J. (2003). How to correct susceptibility distortions in spin-echo echo-planar images: application to diffusion tensor imaging. NeuroImage, 20(2), 870–888.

Andersson, J., Xu, J., Yacoub, E., Auerbach, E., Moeller, S., & Ugurbil, K. (2012). A comprehensive Gaussian process framework for correcting distortions and movements in diffusion images. Proceedings of the 20th Annual Meeting of ISMRM, 2426.

Assaf, Y. (2019). Imaging laminar structures in the gray matter with diffusion MRI. NeuroImage, 197, 677–688.

Assaf, Y., & Pasternak, O. (2008). Diffusion tensor imaging (DTI)-based white matter mapping in brain research: a review. Journal of Molecular Neuroscience: MN, 34(1), 51–61.

Assem, M., Glasser, M. F., Van Essen, D. C., & Duncan, J. (2020). A Domain-General Cognitive Core Defined in Multimodally Parcellated Human Cortex. Cerebral Cortex, 30(8), 4361–4380.

Bartsch, T., & Wulff, P. (2015). The hippocampus in aging and disease: From plasticity to vulnerability. Neuroscience, 309, 1–16.

Bartzokis, G. (2004). Age-related myelin breakdown: a developmental model of cognitive decline and Alzheimer’s disease. Neurobiology of Aging, 25(1), 5–18.

Basser, P. J., & Ozarslan, E. (2010). Anisotropic diffusion: from the apparent diffusion coefficient to the apparent diffusion tensor. Diffusion MRI: Theory, Methods, and Applications. Oxford University Press, Oxford, 79–91.

Bassett, D. S., Cullen, K. E., Eickhoff, S. B., Farah, M. J., Goda, Y., Haggard, P., Hu, H., Hurd, Y. L., Josselyn, S. A., Khakh, B. S., Knoblich, J. A., Poirazi, P., Poldrack, R. A., Prinz, M., Roelfsema, P. R., Spires-Jones, T. L., Sur, M., & Ueda, H. R. (2020). Reflections on the past two decades of neuroscience. Nature Reviews. Neuroscience, 21(10), 524–534.

Bassett, D. S., & Sporns, O. (2017). Network neuroscience. Nature Neuroscience, 20(3), 353–364.

Beaulieu, C. (2002). The basis of anisotropic water diffusion in the nervous system--a technical review. NMR in Biomedicine, 15(7-8), 435–455.

Benedetti, B., Charil, A., Rovaris, M., Judica, E., Valsasina, P., Sormani, M. P., & Filippi, M. (2006). Influence of aging on brain gray and white matter changes assessed by conventional, MT, and DT MRI. Neurology, 66(4), 535–539.

Boutsidis, C., & Gallopoulos, E. (2008). SVD based initialization: A head start for nonnegative matrix factorization. Pattern Recognition, 41(4), 1350–1362.

Bussy, A., Patel, R., Plitman, E., Tullo, S., Salaciak, A., Bedford, S. A., Farzin, S., Béland, M.-L., Valiquette, V., Kazazian, C., Tardif, C. L., Devenyi, G. A., & Chakravarty, M. (2021). Hippocampus shape across the healthy lifespan and its relationship with cognition. Neurobiology of Aging, 106, 153–168.

Bussy, A., Plitman, E., Patel, R., Tullo, S., Salaciak, A., Bedford, S. A., Farzin, S., Béland, M.-L., Valiquette, V., Kazazian, C., Tardif, C. L., Devenyi, G. A., Chakravarty, M. M., & Alzheimer’s Disease Neuroimaging Initiative. (2021). Hippocampal subfield volumes across the healthy lifespan and the effects of MR sequence on estimates. NeuroImage, 233, 117931.

Chad, J. A., Pasternak, O., Salat, D. H., & Chen, J. J. (2018). Re-examining age-related differences in white matter microstructure with free-water corrected diffusion tensor imaging. Neurobiology of Aging, 71, 161–170.

Cox, S. R., Bastin, M. E., Ritchie, S. J., Dickie, D. A., Liewald, D. C., Muñoz Maniega, S., Redmond, P., Royle, N. A., Pattie, A., Valdés Hernández, M., Corley, J., Aribisala, B. S., McIntosh, A. M., Wardlaw, J. M., & Deary, I. J. (2018). Brain cortical characteristics of lifetime cognitive ageing. Brain Structure & Function, 223(1), 509–518.

De Jager, P. L., Shulman, J. M., Chibnik, L. B., Keenan, B. T., Raj, T., Wilson, R. S., Yu, L., Leurgans, S. E., Tran, D., Aubin, C., Anderson, C. D., Biffi, A., Corneveaux, J. J., Huentelman, M. J., Alzheimer’s Disease Neuroimaging Initiative, Rosand, J., Daly, M. J., Myers, A. J., Reiman, E. M., … Evans, D. A. (2012). A genome-wide scan for common variants affecting the rate of age-related cognitive decline. Neurobiology of Aging, 33(5), 1017.e1–e15.

Dickerson, B. C., Feczko, E., Augustinack, J. C., Pacheco, J., Morris, J. C., Fischl, B., & Buckner, R. L. (2009). Differential effects of aging and Alzheimer’s disease on medial temporal lobe cortical thickness and surface area. Neurobiology of Aging, 30(3), 432–440.

Douaud, G., Jbabdi, S., Behrens, T. E. J., Menke, R. A., Gass, A., Monsch, A. U., Rao, A., Whitcher, B., Kindlmann, G., Matthews, P. M., & Smith, S. (2011). DTI measures in crossing-fibre areas: increased diffusion anisotropy reveals early white matter alteration in MCI and mild Alzheimer’s disease. NeuroImage, 55(3), 880–890.

Douaud, G., Menke, R. A. L., Gass, A., Monsch, A. U., Rao, A., Whitcher, B., Zamboni, G., Matthews, P. M., Sollberger, M., & Smith, S. (2013). Brain microstructure reveals early abnormalities more than two years prior to clinical progression from mild cognitive impairment to Alzheimer’s disease. The Journal of Neuroscience: The Official Journal of the Society for Neuroscience, 33(5), 2147–2155.

Edwards, L. J., Kirilina, E., Mohammadi, S., & Weiskopf, N. (2018). Microstructural imaging of human neocortex in vivo. NeuroImage, 182, 184–206.

Esiri, M. M. (2007). Ageing and the brain. The Journal of Pathology, 211(2), 181–187.

Eskildsen, S. F., Coupé, P., Fonov, V., Manjón, J. V., Leung, K. K., Guizard, N., Wassef, S. N., Østergaard, L. R., Collins, D. L., & Alzheimer’s Disease Neuroimaging Initiative. (2012). BEaST: brain extraction based on nonlocal segmentation technique. NeuroImage, 59(3), 2362–2373.

Ewers, M. (2020). Reserve in Alzheimer’s disease: update on the concept, functional mechanisms and sex differences. Current Opinion in Psychiatry, 33(2), 178–184.

Filippini, N., Zsoldos, E., Haapakoski, R., Sexton, C. E., Mahmood, A., Allan, C. L., Topiwala, A., Valkanova, V., Brunner, E. J., Shipley, M. J., Auerbach, E., Moeller, S., Uǧurbil, K., Xu, J., Yacoub, E., Andersson, J., Bijsterbosch, J., Clare, S., Griffanti, L., … Ebmeier, K. P. (2014). Study protocol: The Whitehall II imaging sub-study. BMC Psychiatry, 14, 159.

Fjell, A. M., McEvoy, L., Holland, D., Dale, A. M., Walhovd, K. B., & Alzheimer’s Disease Neuroimaging Initiative. (2014). What is normal in normal aging? Effects of aging, amyloid and Alzheimer’s disease on the cerebral cortex and the hippocampus. Progress in Neurobiology, 117, 20–40.

Fjell, A. M., Westlye, L. T., Grydeland, H., Amlien, I., Espeseth, T., Reinvang, I., Raz, N., Dale, A. M., Walhovd, K. B., & Alzheimer Disease Neuroimaging Initiative. (2014). Accelerating cortical thinning: unique to dementia or universal in aging? Cerebral Cortex, 24(4), 919–934.

Frangou, S., Modabbernia, A., Williams, S. C. R., Papachristou, E., Doucet, G. E., Agartz, I., Aghajani, M., Akudjedu, T. N., Albajes-Eizagirre, A., Alnaes, D., Alpert, K. I., Andersson, M., Andreasen, N. C., Andreassen, O. A., Asherson, P., Banaschewski, T., Bargallo, N., Baumeister, S., Baur-Streubel, R., … Dima, D. (2022). Cortical thickness across the lifespan: Data from 17,075 healthy individuals aged 3-90 years. Human Brain Mapping, 43(1), 431–451.

Glasser, M. F., Coalson, T. S., Robinson, E. C., Hacker, C. D., Harwell, J., Yacoub, E., Ugurbil, K., Andersson, J., Beckmann, C. F., Jenkinson, M., Smith, S. M., & Van Essen, D. C. (2016). A multi-modal parcellation of human cerebral cortex. Nature, 536(7615), 171–178.

Goh, J. O., An, Y., & Resnick, S. M. (2012). Differential trajectories of age-related changes in components of executive and memory processes. Psychology and Aging, 27(3), 707–719.

Goriounova, N. A., Heyer, D. B., Wilbers, R., Verhoog, M. B., Giugliano, M., Verbist, C., Obermayer, J., Kerkhofs, A., Smeding, H., Verberne, M., Idema, S., Baayen, J. C., Pieneman, A. W., de Kock, C. P., Klein, M., & Mansvelder, H. D. (2018). Large and fast human pyramidal neurons associate with intelligence. eLife, 7. https://doi.org/10.7554/eLife.41714

Gregory, S., Long, J. D., Klöppel, S., Razi, A., Scheller, E., Minkova, L., Papoutsi, M., Mills, J. A., Durr, A., Leavitt, B. R., Roos, R. A. C., Stout, J. C., Scahill, R. I., Langbehn, D. R., Tabrizi, S. J., & Rees, G. (2017). Operationalizing compensation over time in neurodegenerative disease. Brain: A Journal of Neurology, 140(4), 1158–1165.

Groves, A. R., Smith, S. M., Fjell, A. M., Tamnes, C. K., Walhovd, K. B., Douaud, G., Woolrich, M. W., & Westlye, L. T. (2012). Benefits of multi-modal fusion analysis on a large-scale dataset: life-span patterns of inter-subject variability in cortical morphometry and white matter microstructure. NeuroImage, 63(1), 365–380.

Grydeland, H., Walhovd, K. B., Tamnes, C. K., Westlye, L. T., & Fjell, A. M. (2013). Intracortical myelin links with performance variability across the human lifespan: results from T1-and T2-weighted MRI myelin mapping and diffusion tensor imaging. The Journal of Neuroscience: The Official Journal of the Society for Neuroscience, 33(47), 18618–18630.

Habeck, C., Gazes, Y., Razlighi, Q., & Stern, Y. (2020). Cortical thickness and its associations with age, total cognition and education across the adult lifespan. PloS One, 15(3), e0230298.

Habes, M., Grothe, M. J., Tunc, B., McMillan, C., Wolk, D. A., & Davatzikos, C. (2020). Disentangling Heterogeneity in Alzheimer’s Disease and Related Dementias Using Data-Driven Methods. Biological Psychiatry. https://doi.org/10.1016/j.biopsych.2020.01.016

Hedman, A. M., van Haren, N. E. M., Schnack, H. G., Kahn, R. S., & Hulshoff Pol, H. E. (2012). Human brain changes across the life span: a review of 56 longitudinal magnetic resonance imaging studies. Human Brain Mapping, 33(8), 1987–2002.

Heim, A. W. (1970). Manual for the AH4 group test of general intelligence. Windsor: NFER.

Helmer, M., Warrington, S. D., & Mohammadi-Nejad, A. R. (2021). On stability of Canonical Correlation Analysis and Partial Least Squares with application to brain-behavior associations. BioRxiv. https://www.biorxiv.org/content/10.1101/2020.08.25.265546v2.abstract

Jagust, W. (2013). Vulnerable neural systems and the borderland of brain aging and neurodegeneration. Neuron, 77(2), 219–234.

Jespersen, S. N., Kroenke, C. D., Østergaard, L., Ackerman, J. J. H., & Yablonskiy, D. A. (2007). Modeling dendrite density from magnetic resonance diffusion measurements. NeuroImage, 34(4), 1473–1486.

Josefsson, M., de Luna, X., Pudas, S., Nilsson, L.-G., & Nyberg, L. (2012). Genetic and lifestyle predictors of 15-year longitudinal change in episodic memory. Journal of the American Geriatrics Society, 60(12), 2308–2312.

Kleinnijenhuis, M., van Mourik, T., Norris, D. G., Ruiter, D. J., van Cappellen van Walsum, A.-M., & Barth, M. (2015). Diffusion tensor characteristics of gyrencephaly using high resolution diffusion MRI in vivo at 7T. NeuroImage, 109, 378–387.

Kochunov, P., Glahn, D. C., Nichols, T. E., Winkler, A. M., Hong, E. L., Holcomb, H. H., Stein, J. L., Thompson, P. M., Curran, J. E., Carless, M. A., Olvera, R. L., Johnson, M. P., Cole, S. A., Kochunov, V., Kent, J., & Blangero, J. (2011). Genetic analysis of cortical thickness and fractional anisotropy of water diffusion in the brain. Frontiers in Neuroscience, 5, 120.

Krishnan, A., Williams, L. J., McIntosh, A. R., & Abdi, H. (2011). Partial Least Squares (PLS) methods for neuroimaging: a tutorial and review. NeuroImage, 56(2), 455–475.

Lampinen, B., Szczepankiewicz, F., Novén, M., van Westen, D., Hansson, O., Englund, E., Mårtensson, J., Westin, C.-F., & Nilsson, M. (2019). Searching for the neurite density with diffusion MRI: Challenges for biophysical modeling. Human Brain Mapping, 40(8), 2529–2545.

Lazari, A., & Lipp, I. (2021). Can MRI measure myelin? Systematic review, qualitative assessment, and meta-analysis of studies validating microstructural imaging with myelin histology. NeuroImage, 230, 117744.

Lebel, C., Gee, M., Camicioli, R., Wieler, M., Martin, W., & Beaulieu, C. (2012). Diffusion tensor imaging of white matter tract evolution over the lifespan. NeuroImage, 60(1), 340–352.

Le Bihan, D., Mangin, J. F., Poupon, C., Clark, C. A., Pappata, S., Molko, N., & Chabriat, H. (2001). Diffusion tensor imaging: concepts and applications. Journal of Magnetic Resonance imaging: JMRI, 13(4), 534–546.

Lee, D. D., & Seung, H. S. (1999). Learning the parts of objects by non-negative matrix factorization. Nature, 401(6755), 788–791.

Lee, P., Kim, H.-R., Jeong, Y., & Alzheimer’s Disease Neuroimaging Initiative. (2020). Detection of gray matter microstructural changes in Alzheimer’s disease continuum using fiber orientation. BMC Neurology, 20(1), 362.

Lemaitre, H., Goldman, A. L., Sambataro, F., Verchinski, B. A., Meyer-Lindenberg, A., Weinberger, D. R., & Mattay, V. S. (2012). Normal age-related brain morphometric changes: nonuniformity across cortical thickness, surface area and gray matter volume? Neurobiology of Aging, 33(3), 617.e1–e9.

Lerch, J. P., & Evans, A. C. (2005). Cortical thickness analysis examined through power analysis and a population simulation. NeuroImage, 24(1), 163–173.

Lerch, J. P., van der Kouwe, A. J. W., Raznahan, A., Paus, T., Johansen-Berg, H., Miller, K. L., Smith, S. M., Fischl, B., & Sotiropoulos, S. N. (2017). Studying neuroanatomy using MRI. Nature Neuroscience, 20(3), 314–326.

Leuze, C. W. U., Anwander, A., Bazin, P.-L., Dhital, B., Stüber, C., Reimann, K., Geyer, S., & Turner, R. (2014). Layer-specific intracortical connectivity revealed with diffusion MRI. Cerebral Cortex, 24(2), 328–339.

Loken, C., Gruner, D., Groer, L., Peltier, R., Bunn, N., Craig, M., Henriques, T., Dempsey, J., Yu, C.-H., Chen, J., Jonathan Dursi, L., Chong, J., Northrup, S., Pinto, J., Knecht, N., & Van Zon, R. (2010). SciNet: Lessons Learned from Building a Power-efficient Top-20 System and Data Centre. Journal of Physics. Conference Series, 256(1), 012026.

Lowe, A. J., Paquola, C., Vos de Wael, R., Girn, M., Lariviere, S., Tavakol, S., Caldairou, B., Royer, J., Schrader, D. V., Bernasconi, A., Bernasconi, N., Spreng, R. N., & Bernhardt, B. C. (2019). Targeting age-related differences in brain and cognition with multimodal imaging and connectome topography profiling. Human Brain Mapping, 40(18), 5213–5230.

Madden, D. J., Bennett, I. J., Burzynska, A., Potter, G. G., Chen, N.-K., & Song, A. W. (2012). Diffusion tensor imaging of cerebral white matter integrity in cognitive aging. Biochimica et Biophysica Acta, 1822(3), 386–400.

Mancini, M., Karakuzu, A., Cohen-Adad, J., Cercignani, M., Nichols, T. E., & Stikov, N. (2020). An interactive meta-analysis of MRI biomarkers of myelin. eLife, 9. https://doi.org/10.7554/eLife.61523

Manjón, J. V., Coupé, P., Martí-Bonmatí, L., Collins, D. L., & Robles, M. (2010). Adaptive non-local means denoising of MR images with spatially varying noise levels. Journal of Magnetic Resonance Imaging: JMRI, 31(1), 192–203.

Marek, S., Tervo-Clemmens, B., Calabro, F. J., Montez, D. F., Kay, B. P., Hatoum, A. S., Donohue, M. R., Foran, W., Miller, R. L., Feczko, E., Miranda-Dominguez, O., Graham, A. M., Earl, E. A., Perrone, A. J., Cordova, M., Doyle, O., Moore, L. A., Conan, G., Uriarte, J., … Dosenbach, N. U. F. (2020). Towards Reproducible Brain-Wide Association Studies. In bioRxiv (p. 2020.08.21.257758). https://doi.org/10.1101/2020.08.21.257758

Marmot, M., & Brunner, E. (2005). Cohort Profile: the Whitehall II study. International Journal of Epidemiology, 34(2), 251–256.

Marner, L., Nyengaard, J. R., Tang, Y., & Pakkenberg, B. (2003). Marked loss of myelinated nerve fibers in the human brain with age. The Journal of Comparative Neurology, 462(2), 144–152.

McIntosh, A. R., & Lobaugh, N. J. (2004). Partial least squares analysis of neuroimaging data: applications and advances. NeuroImage, 23 Suppl 1, S250–S263.

McIntosh, A. R., & Mišić, B. (2013). Multivariate statistical analyses for neuroimaging data. Annual Review of Psychology, 64, 499–525.

McKavanagh, R., Torso, M., Jenkinson, M., Kolasinski, J., Stagg, C. J., Esiri, M. M., McNab, J. A., Johansen-Berg, H., Miller, K. L., & Chance, S. A. (2019). Relating diffusion tensor imaging measurements to microstructural quantities in the cerebral cortex in multiple sclerosis. Human Brain Mapping, 40(15), 4417–4431.

McNab, J. A., Polimeni, J. R., Wang, R., Augustinack, J. C., Fujimoto, K., Stevens, A., Triantafyllou, C., Janssens, T., Farivar, R., Folkerth, R. D., Vanduffel, W., & Wald, L. L. (2013). Surface based analysis of diffusion orientation for identifying architectonic domains in the in vivo human cortex. NeuroImage, 69, 87–100.

Miller, K. L., Alfaro-Almagro, F., Bangerter, N. K., Thomas, D. L., Yacoub, E., Xu, J., Bartsch, A. J., Jbabdi, S., Sotiropoulos, S. N., Andersson, J. L. R., Griffanti, L., Douaud, G., Okell, T. W., Weale, P., Dragonu, I., Garratt, S., Hudson, S., Collins, R., Jenkinson, M., … Smith, S. M. (2016). Multimodal population brain imaging in the UK Biobank prospective epidemiological study. Nature Neuroscience, 19(11), 1523–1536.

Mole, J., Foley, J., Shallice, T., & Cipolotti, L. (2021). The left frontal lobe is critical for the AH4 fluid intelligence test. Intelligence, 87, 101564.

Nair, G., Tanahashi, Y., Low, H. P., Billings-Gagliardi, S., Schwartz, W. J., & Duong, T. Q. (2005). Myelination and long diffusion times alter diffusion-tensor-imaging contrast in myelin-deficient shiverer mice. NeuroImage, 28(1), 165–174.

Nassar, R., Kaczkurkin, A. N., Xia, C. H., Sotiras, A., Pehlivanova, M., Moore, T. M., Garcia de La Garza, A., Roalf, D. R., Rosen, A. F. G., Lorch, S. A., Ruparel, K., Shinohara, R. T., Davatzikos, C., Gur, R. C., Gur, R. E., & Satterthwaite, T. D. (2018). Gestational Age is Dimensionally Associated with Structural Brain Network Abnormalities Across Development. Cerebral Cortex. https://doi.org/10.1093/cercor/bhy091

Nordin, K., Persson, J., Stening, E., Herlitz, A., Larsson, E.-M., & Söderlund, H. (2018). Structural whole-brain covariance of the anterior and posterior hippocampus: Associations with age and memory. Hippocampus, 28(2), 151–163.

Oschwald, J., Guye, S., Liem, F., Rast, P., Willis, S., Röcke, C., Jäncke, L., Martin, M., & Mérillat, S. (2019). Brain structure and cognitive ability in healthy aging: a review on longitudinal correlated change. Reviews in the Neurosciences, 31(1), 1–57.

Panizzon, M. S., Fennema-Notestine, C., Eyler, L. T., Jernigan, T. L., Prom-Wormley, E., Neale, M., Jacobson, K., Lyons, M. J., Grant, M. D., Franz, C. E., Xian, H., Tsuang, M., Fischl, B., Seidman, L., Dale, A., & Kremen, W. S. (2009). Distinct genetic influences on cortical surface area and cortical thickness. Cerebral Cortex, 19(11), 2728–2735.

Park, D. C., & Reuter-Lorenz, P. (2009). The Adaptive Brain: Aging and Neurocognitive Scaffolding. Annual Review of Psychology, 60(1), 173–196.

Patel, R., Steele, C. J., Chen, A. G. X., Patel, S., Devenyi, G. A., Germann, J., Tardif, C. L., & Chakravarty, M. M. (2020). Investigating microstructural variation in the human hippocampus using non-negative matrix factorization. NeuroImage, 207, 116348.

Patel, Y., Parker, N., Shin, J., Howard, D., French, L., Thomopoulos, S. I., Pozzi, E., Abe, Y., Abé, C., Anticevic, A., & Others. (2021). Virtual histology of cortical thickness and shared neurobiology in 6 psychiatric disorders. JAMA Psychiatry, 78(1), 47–63.

Persson, J., Nyberg, L., Lind, J., Larsson, A., Nilsson, L.-G., Ingvar, M., & Buckner, R. L. (2006). Structure-function correlates of cognitive decline in aging. Cerebral Cortex, 16(7), 907–915.

Persson, J., Pudas, S., Lind, J., Kauppi, K., Nilsson, L.-G., & Nyberg, L. (2012). Longitudinal structure-function correlates in elderly reveal MTL dysfunction with cognitive decline. Cerebral Cortex, 22(10), 2297–2304.

Philippi, N., Noblet, V., Duron, E., Cretin, B., Boully, C., Wisniewski, I., Seux, M. L., Martin-Hunyadi, C., Chaussade, E., Demuynck, C., Kremer, S., Lehéricy, S., Gounot, D., Armspach, J. P., Hanon, O., & Blanc, F. (2016). Exploring anterograde memory: a volumetric MRI study in patients with mild cognitive impairment. Alzheimer’s Research & Therapy, 8(1), 26.

Pichet Binette, A., Gonneaud, J., Vogel, J. W., La Joie, R., Rosa-Neto, P., Collins, D. L., Poirier, J., Breitner, J. C. S., Villeneuve, S., Vachon-Presseau, E., Alzheimer’s Disease Neuroimaging Initiative, & PREVENT-AD Research Group. (2020). Morphometric network differences in ageing versus Alzheimer’s disease dementia. Brain: A Journal of Neurology, 143(2), 635–649.

Ponce, M., van Zon, R., Northrup, S., Gruner, D., Chen, J., Ertinaz, F., Fedoseev, A., Groer, L., Mao, F., Mundim, B. C., Nolta, M., Pinto, J., Saldarriaga, M., Slavnic, V., Spence, E., Yu, C.-H., & Peltier, W. R. (2019). Deploying a Top-100 Supercomputer for Large Parallel Workloads: the Niagara Supercomputer. Proceedings of the Practice and Experience in Advanced Research Computing on Rise of the Machines (learning), 1–8.

Preziosa, P., Kiljan, S., Steenwijk, M. D., Meani, A., van de Berg, W. D. J., Schenk, G. J., Rocca, M. A., Filippi, M., Geurts, J. J. G., & Jonkman, L. E. (2019). Axonal degeneration as substrate of fractional anisotropy abnormalities in multiple sclerosis cortex. Brain: A Journal of Neurology, 142(7), 1921–1937.

Pudas, S., Persson, J., Josefsson, M., de Luna, X., Nilsson, L.-G., & Nyberg, L. (2013). Brain characteristics of individuals resisting age-related cognitive decline over two decades. The Journal of Neuroscience: The Official Journal of the Society for Neuroscience, 33(20), 8668–8677.

Querbes, O., Aubry, F., Pariente, J., Lotterie, J.-A., Démonet, J.-F., Duret, V., Puel, M., Berry, I., Fort, J.-C., Celsis, P., & Alzheimer’s Disease Neuroimaging Initiative. (2009). Early diagnosis of Alzheimer’s disease using cortical thickness: impact of cognitive reserve. Brain: A Journal of Neurology, 132(Pt 8), 2036–2047.

Rakic, P. (1988). Specification of cerebral cortical areas. Science, 241(4862), 170–176.

Rakic, P. (2000). Radial unit hypothesis of neocortical expansion. Novartis Foundation Symposium, 228, 30–42; discussion 42–52.

Raven, J. C. (1958). Guide to using the Mill Hill Vocabulary Scale with the Progressive Matrices Scales. 64. https://psycnet.apa.org/fulltext/1960-00086-000.pdf

Raz, N., & Rodrigue, K. M. (2006). Differential aging of the brain: Patterns, cognitive correlates and modifiers. Neuroscience and Biobehavioral Reviews, 30(6), 730–748.

Reas, E. T., Hagler, D. J., Jr, White, N. S., Kuperman, J. M., Bartsch, H., Wierenga, C. E., Galasko, D., Brewer, J. B., Dale, A. M., & McEvoy, L. K. (2018). Microstructural brain changes track cognitive decline in mild cognitive impairment. NeuroImage. Clinical, 20, 883–891.

Robert, C., Patel, R., Blostein, N., Steele, C. J., & Chakravarty, M. M. (2022). Analyses of microstructural variation in the human striatum using non-negative matrix factorization. NeuroImage, 246, 118744.

Rodríguez-Aranda, C., Waterloo, K., Johnsen, S. H., Eldevik, P., Sparr, S., Wikran, G. C., Herder, M., & Vangberg, T. R. (2016). Neuroanatomical correlates of verbal fluency in early Alzheimer’s disease and normal aging. Brain and Language, 155-156, 24–35.

Rönnlund, M., Nyberg, L., Bäckman, L., & Nilsson, L.-G. (2005). Stability, growth, and decline in adult life span development of declarative memory: cross-sectional and longitudinal data from a population-based study. Psychology and Aging, 20(1), 3–18.

Salat, D. H., Buckner, R. L., Snyder, A. Z., Greve, D. N., Desikan, R. S. R., Busa, E., Morris, J. C., Dale, A. M., & Fischl, B. (2004). Thinning of the cerebral cortex in aging. Cerebral Cortex, 14(7), 721–730.

Salthouse, T. (2010). Selective review of cognitive aging. Journal of International Neuropsychology, 16(5), 754–760.

Schneider, A. L. C., Senjem, M. L., Wu, A., Gross, A., Knopman, D. S., Gunter, J. L., Schwarz, C. G., Mosley, T. H., Gottesman, R. F., Sharrett, A. R., & Jack, C. R., Jr. (2019). Neural correlates of domain-specific cognitive decline: The ARIC-NCS Study. Neurology, 92(10), e1051–e1063.

Scola, E., Bozzali, M., Agosta, F., Magnani, G., Franceschi, M., Sormani, M. P., Cercignani, M., Pagani, E., Falautano, M., Filippi, M., & Falini, A. (2010). A diffusion tensor MRI study of patients with MCI and AD with a 2-year clinical follow-up. Journal of Neurology, Neurosurgery, and Psychiatry, 81(7), 798–805.

Seidlitz, J., Váša, F., Shinn, M., Romero-Garcia, R., Whitaker, K. J., Vértes, P. E., Wagstyl, K., Kirkpatrick Reardon, P., Clasen, L., Liu, S., Messinger, A., Leopold, D. A., Fonagy, P., Dolan, R. J., Jones, P. B., Goodyer, I. M., NSPN Consortium, Raznahan, A., & Bullmore, E. T. (2018). Morphometric Similarity Networks Detect Microscale Cortical Organization and Predict Inter-Individual Cognitive Variation. Neuron, 97(1), 231–247.e7.

Shaw, M. E., Sachdev, P. S., Anstey, K. J., & Cherbuin, N. (2016). Age-related cortical thinning in cognitively healthy individuals in their 60s: the PATH Through Life study. Neurobiology of Aging, 39, 202–209.

Singh-Manoux, A., Kivimaki, M., Glymour, M. M., Elbaz, A., Berr, C., Ebmeier, K. P., Ferrie, J. E., & Dugravot, A. (2012). Timing of onset of cognitive decline: results from Whitehall II prospective cohort study. BMJ, 344, d7622.

Soldan, A., Pettigrew, C., & Albert, M. (2020). Cognitive Reserve from the Perspective of Preclinical Alzheimer Disease: 2020 Update. Clinics in Geriatric Medicine, 36(2), 247–263.

Song, S.-K., Kim, J. H., Lin, S.-J., Brendza, R. P., & Holtzman, D. M. (2004). Diffusion tensor imaging detects age-dependent white matter changes in a transgenic mouse model with amyloid deposition. Neurobiology of Disease, 15(3), 640–647.

Song, S.-K., Sun, S.-W., Ramsbottom, M. J., Chang, C., Russell, J., & Cross, A. H. (2002). Dysmyelination revealed through MRI as increased radial (but unchanged axial) diffusion of water. NeuroImage, 17(3), 1429–1436.

Sotiras, A., Resnick, S. M., & Davatzikos, C. (2015). Finding imaging patterns of structural covariance via Non-Negative Matrix Factorization. NeuroImage, 108, 1–16.

Sotiras, A., Toledo, J. B., Gur, R. E., Gur, R. C., Satterthwaite, T. D., & Davatzikos, C. (2017). Patterns of coordinated cortical remodeling during adolescence and their associations with functional specialization and evolutionary expansion. Proceedings of the National Academy of Sciences of the United States of America, 114(13), 3527–3532.

Sotiropoulos, S. N., Jbabdi, S., Xu, J., Andersson, J. L., Moeller, S., Auerbach, E. J., Glasser, M. F., Hernandez, M., Sapiro, G., Jenkinson, M., Feinberg, D. A., Yacoub, E., Lenglet, C., Van Essen, D. C., Ugurbil, K., Behrens, T. E. J., & WU-Minn HCP Consortium. (2013). Advances in diffusion MRI acquisition and processing in the Human Connectome Project. NeuroImage, 80, 125–143.

Stern, Y., Arenaza-Urquijo, E. M., Bartrés-Faz, D., Belleville, S., Cantilon, M., Chetelat, G., Ewers, M., Franzmeier, N., Kempermann, G., Kremen, W. S., Okonkwo, O., Scarmeas, N., Soldan, A., Udeh-Momoh, C., Valenzuela, M., Vemuri, P., Vuoksimaa, E., and the Reserve, & Resilience and Protective Factors PIA Empirical Definitions and Conceptual Frameworks Workgroup. (2020). Whitepaper: Defining and investigating cognitive reserve, brain reserve, and brain maintenance. In Alzheimer’s & Dementia (Vol. 16, Issue 9, pp. 1305–1311). https://doi.org/10.1016/j.jalz.2018.07.219

Storsve, A. B., Fjell, A. M., Tamnes, C. K., Westlye, L. T., Overbye, K., Aasland, H. W., & Walhovd, K. B. (2014). Differential longitudinal changes in cortical thickness, surface area and volume across the adult life span: regions of accelerating and decelerating change. The Journal of Neuroscience: The Official Journal of the Society for Neuroscience, 34(25), 8488–8498.

Sun, S.-W., Liang, H.-F., Cross, A. H., & Song, S.-K. (2008). Evolving Wallerian degeneration after transient retinal ischemia in mice characterized by diffusion tensor imaging. NeuroImage, 40(1), 1–10.

Suri, S., Topiwala, A., Chappell, M. A., Okell, T. W., Zsoldos, E., Singh-Manoux, A., Kivimäki, M., Mackay, C. E., & Ebmeier, K. P. (2019). Association of Midlife Cardiovascular Risk Profiles With Cerebral Perfusion at Older Ages. JAMA Network Open, 2(6), e195776.

Tamnes, C. K., Walhovd, K. B., Dale, A. M., Østby, Y., Grydeland, H., Richardson, G., Westlye, L. T., Roddey, J. C., Hagler, D. J., Due-Tønnessen, P., Holland, D., & Fjell, A. M. (2013). Brain development and aging: Overlapping and unique patterns of change. NeuroImage, 68, 63–74.

Tardif, C. L., Gauthier, C. J., Steele, C. J., Bazin, P.-L., Schäfer, A., Schaefer, A., Turner, R., & Villringer, A. (2016). Advanced MRI techniques to improve our understanding of experience-induced neuroplasticity. NeuroImage, 131, 55–72.

Torso, M., Bozzali, M., Zamboni, G., Jenkinson, M., Chance, S. A., & Alzheimers Disease Neuroimage Initiative. (2021). Detection of Alzheimer’s Disease using cortical diffusion tensor imaging. Human Brain Mapping, 42(4), 967–977.

Torso, M., Ridgway, G. R., Jenkinson, M., Chance, S., & Frontotemporal Lobar Degeneration Neuroimaging Initiative and the 4-Repeat Tau Neuroimaging Initiative (4RTNI). (2021). Intracortical diffusion tensor imaging signature of microstructural changes in frontotemporal lobar degeneration. Alzheimer’s Research & Therapy, 13(1), 180.

Truong, T.-K., Guidon, A., & Song, A. W. (2014). Cortical depth dependence of the diffusion anisotropy in the human cortical gray matter in vivo. PloS One, 9(3), e91424.

Tucker-Drob, E. M. (2011a). Global and domain-specific changes in cognition throughout adulthood. Developmental Psychology, 47(2), 331–343.

Tucker-Drob, E. M. (2011b). Neurocognitive functions and everyday functions change together in old age. Neuropsychology, 25(3), 368–377.

Tucker-Drob, E. M., & Salthouse, T. A. (2011). Individual Differences in Cognitive Aging. In T. Chamorro-Premuzic, S. von Stumm, & A. Furnham (Eds.), The Wiley-Blackwell Handbook of Individual Differences (Vol. 132, pp. 242–267). Wiley-Blackwell.

Tullo, S., Patel, R., Devenyi, G. A., Salaciak, A., Bedford, S. A., Farzin, S., Wlodarski, N., Tardif, C. L., Group, P.-A. R., Breitner, J. C. S., & Others. (2019). MR-based age-related effects on the striatum, globus pallidus, and thalamus in healthy individuals across the adult lifespan. Human Brain Mapping, 40(18), 5269–5288.

Tuor, U. I., Morgunov, M., Sule, M., Qiao, M., Clark, D., Rushforth, D., Foniok, T., & Kirton, A. (2014). Cellular correlates of longitudinal diffusion tensor imaging of axonal degeneration following hypoxic-ischemic cerebral infarction in neonatal rats. NeuroImage. Clinical, 6, 32–42.

Tustison, N. J., Avants, B. B., Cook, P. A., Zheng, Y., Egan, A., Yushkevich, P. A., & Gee, J. C. (2010). N4ITK: improved N3 bias correction. IEEE Transactions on Medical Imaging, 29(6), 1310–1320.

Uddin, M. N., Figley, T. D., Solar, K. G., Shatil, A. S., & Figley, C. R. (2019). Comparisons between multi-component myelin water fraction, T1w/T2w ratio, and diffusion tensor imaging measures in healthy human brain structures. Scientific Reports, 9(1), 2500.

van den Heuvel, M. P., & Sporns, O. (2013). Network hubs in the human brain. Trends in Cognitive Sciences, 17(12), 683–696.

Varikuti, D. P., Genon, S., Sotiras, A., Schwender, H., Hoffstaedter, F., Patil, K. R., Jockwitz, C., Caspers, S., Moebus, S., Amunts, K., Davatzikos, C., & Eickhoff, S. B. (2018). Evaluation of non-negative matrix factorization of grey matter in age prediction. NeuroImage, 173, 394–410.

Vidal-Pineiro, D., Parker, N., Shin, J., French, L., Grydeland, H., Jackowski, A. P., Mowinckel, A. M., Patel, Y., Pausova, Z., Salum, G., Sørensen, Ø., Walhovd, K. B., Paus, T., Fjell, A. M., & Alzheimer’s Disease Neuroimaging Initiative and the Australian Imaging Biomarkers and Lifestyle flagship study of ageing. (2020). Cellular correlates of cortical thinning throughout the lifespan. Scientific Reports, 10(1), 21803.

Walhovd, K. B., Krogsrud, S. K., Amlien, I. K., Bartsch, H., Bjørnerud, A., Due-Tønnessen, P., Grydeland, H., Hagler, D. J., Jr, Håberg, A. K., Kremen, W. S., Ferschmann, L., Nyberg, L., Panizzon, M. S., Rohani, D. A., Skranes, J., Storsve, A. B., Sølsnes, A. E., Tamnes, C. K., Thompson, W. K., … Fjell, A. M. (2016). Neurodevelopmental origins of lifespan changes in brain and cognition. Proceedings of the National Academy of Sciences of the United States of America, 113(33), 9357–9362.

Walhovd, K. B., Westlye, L. T., Amlien, I., Espeseth, T., Reinvang, I., Raz, N., Agartz, I., Salat, D. H., Greve, D. N., Fischl, B., Dale, A. M., & Fjell, A. M. (2011). Consistent neuroanatomical age-related volume differences across multiple samples. Neurobiology of Aging, 32(5), 916–932.

Wilson, R. S., Beckett, L. A., Barnes, L. L., Schneider, J. A., Bach, J., Evans, D. A., & Bennett, D. A. (2002). Individual differences in rates of change in cognitive abilities of older persons. Psychology and Aging, 17(2), 179–193.

Zatorre, R. J., Fields, R. D., & Johansen-Berg, H. (2012). Plasticity in gray and white: neuroimaging changes in brain structure during learning. Nature Neuroscience, 15(4), 528–536.

Zeighami, Y., Fereshtehnejad, S.-M., Dadar, M., Collins, D. L., Postuma, R. B., Mišić, B., & Dagher, A. (2017). A clinical-anatomical signature of Parkinson’s disease identified with partial least squares and magnetic resonance imaging. NeuroImage. https://doi.org/10.1016/j.neuroimage.2017.12.050

Zhang, H., Sachdev, P. S., Wen, W., Kochan, N. A., Crawford, J. D., Brodaty, H., Slavin, M. J., Reppermund, S., Kang, K., & Trollor, J. N. (2013). Grey matter correlates of three language tests in non-demented older adults. PloS One, 8(11), e80215.

