## Supplementary Material for "Individual variation in brain structural-cognition relationships in aging"

### Supplementary Information

#### Sample Selection

Of the 800 individuals randomly selected to participate in the Whitehall II Imaging Sub-Study, 25 participants did not receive an MRI scan. Of the 775 with an available MRI scan, 32 subjects were excluded due to incidental findings such as large strokes, brain tumours, or cysts. Furthermore, 243 were excluded due to incomplete whole-brain coverage in the diffusion weighted imaging acquisition (incomplete coverage of the cerebellum and occipital lobe). Motion quality control was assessed for each of the remaining 500 participants according to in-lab protocols (https://github.com/CoBrALab/documentation/wiki/Motion-Quality-Control-Manual), with 31 participants failing motion quality control. A further 48 participants were excluded as they failed CIVET processing quality control with failure modes including incorrect tissue classification, failed registration, and incorrect white surface extraction (typically characterized by bridging artifacts as described in http://www.bic.mni.mcgill.ca/ServicesSoftware/CIVET-2-1-0-Quality-Control). Of the 421 participants with whole brain diffusion weighted imaging coverage and who passed motion and civet quality control, a further 23 were excluded due to incomplete longitudinal cognitive assessments resulting in a final analysis sample of 398 individuals. None of the participants had a clinical diagnosis of dementia at the time of the MRI scan (Figure S1).

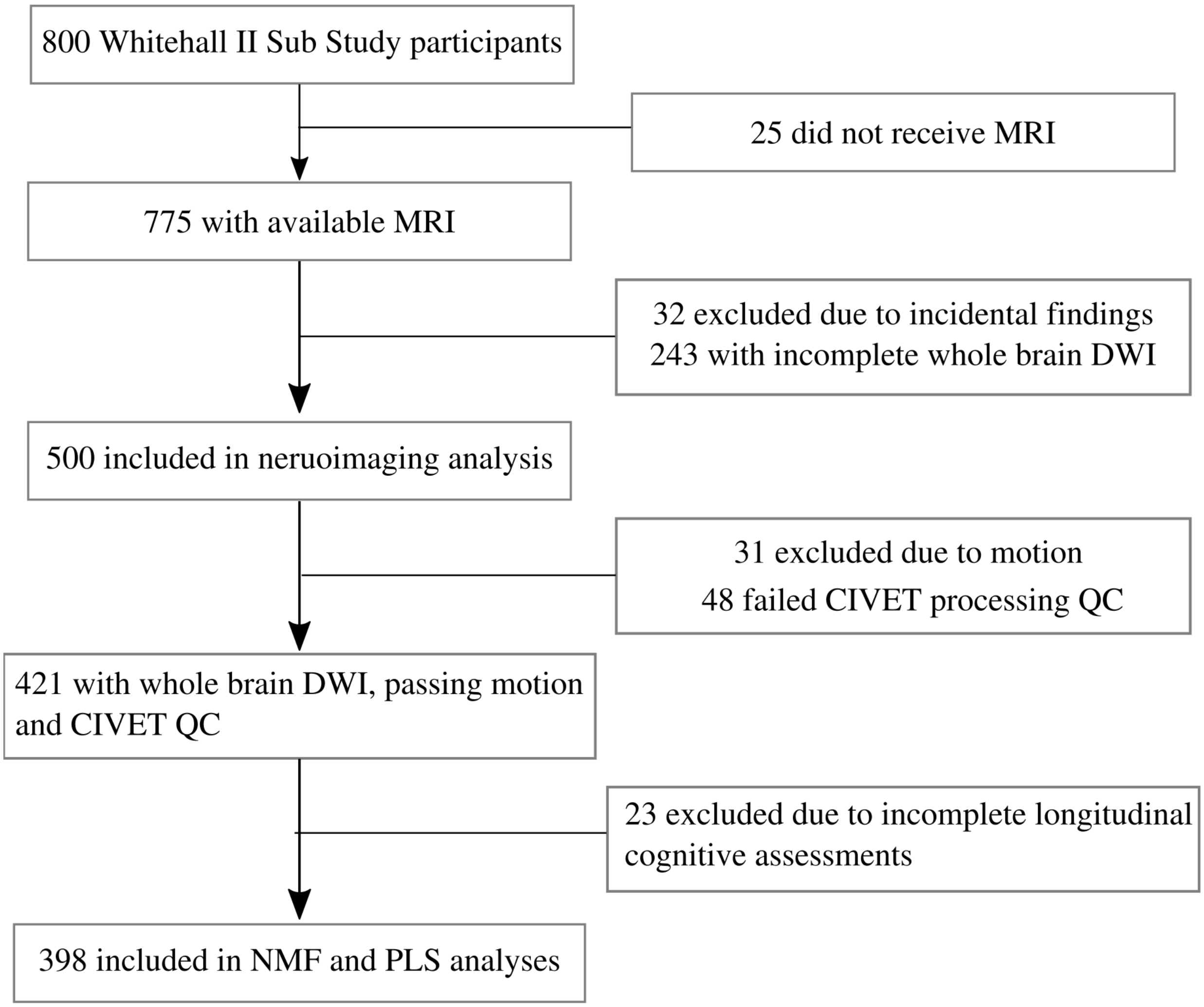


**Figure S1:** Flowchart of exclusion process to arrive at analysis sample of 398 individuals

#### Split half stability analysis

Split half stability analysis was used to identify 10 components as a suitable balance between spatial stability and reconstruction accuracy (Figure S2). Figure S2 plots the stability coefficient at each granularity analyzed, describing the correlation of vertex component values in split half datasets. A high stability coefficient is achieved when the output components consistently represent the entire sample as opposed to being driven by a smaller subset of outliers. Stability is highest when the number of components (k) is low; it was moderate but variable at k=2, then peaked in the range of k=4 to k=8, before experiencing a moderate decrease to k=10, a large decrease to k=12, followed by consistent decrease up until a plateauing around k=60. These results suggest the range of k=4-10 would suitably identify a stable parcellation.

We also examined the gradient of the reconstruction error, representing the gain in accuracy achieved by increasing the number of components (Figure S2, blue). As expected, a finer parcellation (more components) resulted in higher accuracy. However, the magnitude of these gains were notably larger with fewer components, showing that the gain in accuracy of the NMF model at high granularities is much lower. Up until k=10, increasing the number of components resulted in large gains in accuracy, suggesting that k>=10 was a suitable choice for capturing the major patterns in the brain matrix. Together with the stability results, we selected k=10 for further analysis.

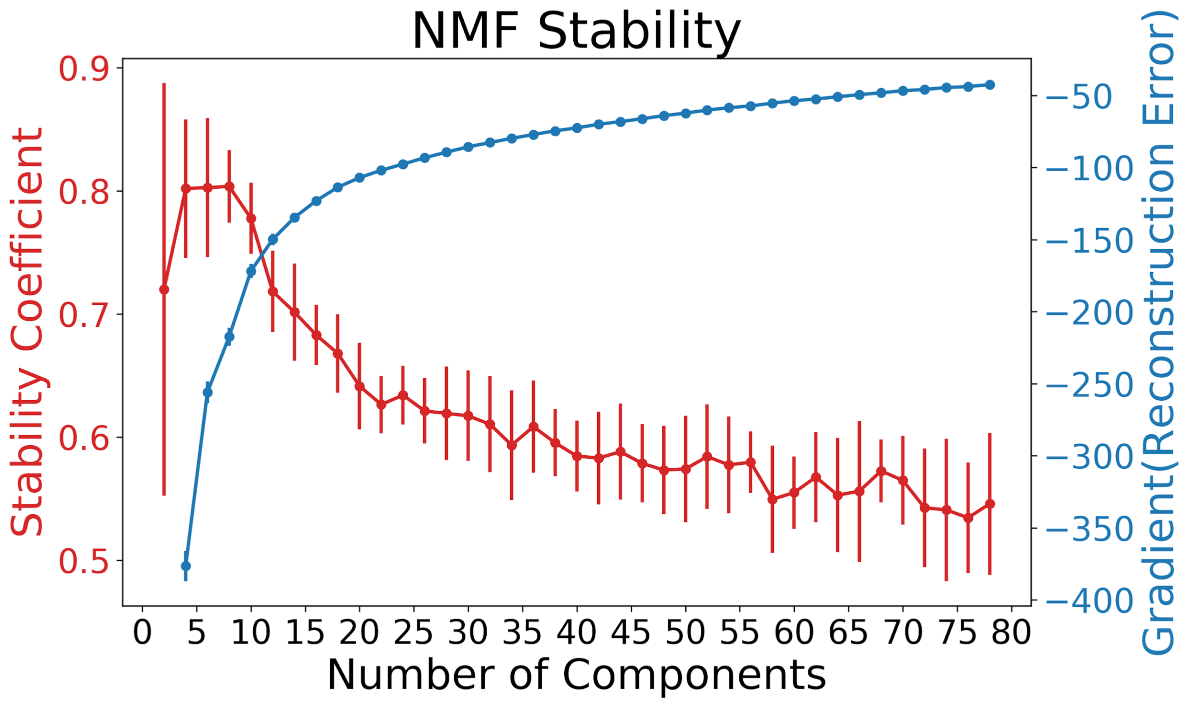


**Figure S2.** Split half stability coefficient and change in reconstruction errors for 2-80 component decomposition. To balance high stability (red line) while capturing major changes in accuracy (blue), 10 components was selected for further analysis.

#### Longitudinal Cognitive Performance

Semantic fluency, inductive, verbal, and mathematical reasoning, and vocabulary performance declined significantly over the 30-year follow-up from Waves 5 to 11 (summary statistics in Table S1). Lexical fluency remained stable and short-term memory showed a trend to decline (Table S1).

**Table S1:** Summary statistics of the linear mixed effects model for cognitive change from Wave 5 to Wave 11 of the Whitehall II Study. Unstandardized beta coefficients and uncorrected p-values for each of the time since baseline, baseline age, and interaction effects for each cognitive test. Using a bonferroni threshold of p<0.05/7 (p<0.00714), significant associations are bolded and italicized.

| | Time | | Baseline Age | | Time*Baseline Age | |
|---|---|---|---|---|---|---|
| Test | B [95% CI] | p | B [95% CI] | p | B [95% CI] | p |
| Lexical Fluency | -0.08[-0.13,-0.04] | **0.0002** | -0.1 [-0.18,-0.02] | 0.01 | -0.003 [-0.01,0.001] | 0.17 |
| Semantic Fluency | -0.06[-0.1,-0.02] | **0.007** | -0.09 [-0.16,-0.01] | 0.0268 | -0.005 [-0.01,-0.001] | 0.0199 |
| Short Term Memory | -0.03[-0.06,0.003] | 0.08 | -0.07 [-0.11,-0.03] | **0.0008** | -0.003 [-0.006,0.0001] | 0.0509 |
| Inductive Reasoning | -0.06[-0.12,0.004] | *0.069* | -0.23 [-0.4,-0.05] | 0.01 | -0.019 [-0.025,-0.12] | ***0*** |
| Verbal Reasoning | -0.03 [-0.06,0.005] | *0.096* | -0.1 [-0.19,-0.02] | 0.0175 | -0.01 [-0.014,-0.006] | ***0*** |
| Mathematical Reasoning | -0.03 [-0.07,0.01] | *0.17* | -0.13 [-0.23,-0.02] | 0.0148 | -0.009 [-0.013,-0.005] | ***0*** |
| Vocabulary | 0.05[0.02,0.08] | ***0.0006*** | 0.04[-0.03,0.12] | 0.29 | -0.005 [-0.008,-0.002] | ***0.0009*** |

### Predicting Future Cognition Using Brain Cognition Patterns

To further investigate the utility and impact of PLS derived LVs, we performed linear models to examine the relationship between LV behaviour scores and Wave 12 cognitive performance. Results of this analysis are in section 2.4 of the main text. Here we provide supplementary results on the demographic characterization of LV scores. We assessed sex differences in LV scores using Welch's two sample t-tests to detect differences in mean LV scores between males and females. Both LV1 and LV2 showed significant sex differences with females having higher LV1 (t(123.9)=4.06, p=0.00008477) but lower LV2 (t(152.53)=-3.79,p=0.0002161) (Figure S3). We assessed the relationship between age and LV scores using a linear model with age as dependent variable and LV behaviour score as the independent variable with no other covariates. A separate model was run for each of LV1 and LV2, with neither showing significant relationships (LV1: beta=-0.03, p=0.697; LV2: beta=0.07, p=0.722) (Figure S3).

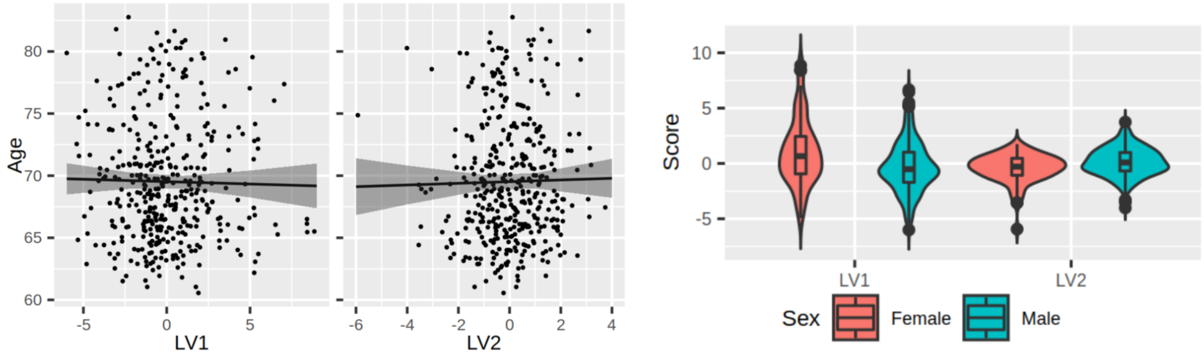


**Fig. S3.** Demographic comparisons of the LV behaviour scores. LV1 and LV2 showed no relation to age (left) but did show sex differences (right).

*In section 2.4 of the main text we present results of linear models assessing the relationship between LV scores and wave 12 performance. In Table S2 below we present these results while controlling for the time interval between the final timepoints which was variable across subjects. This assesses how the future prediction results are impacted by how far ahead these predictions were made. Table S2 is analogous to Table 3 of the main text, but includes the time interval as a covariate. Negligible differences confirm the time interval did not play a role in the relationship between LV scores and future performance.*

***Table S2:*** *Statistical results from linear models run for Wave 12 performance on each of 6 cognitive tests, while covarying for the time between the imaging timepoint and wave 12 which varied across individuals. Each model had the form* $Test \sim LV1 + LV2 + Age + Sex + Education + TimeGap$*, where TimeGap represents the time in years between the imaging timepoint and wave 12. Standardized beta coefficients and 95% confidence intervals are reported, where the standardized coefficient represents the degree of change in wave 12 test scores for every one unit increase in LV scores, in units of standard deviation. Bold p-values are deemed significant at a Bonferroni-corrected threshold of p<0.0083.*

| ***Test*** | ***Term*** | ***Standardized β [95% CI]*** | ***p*** |
|---|---|---|---|
| *Semantic Fluency* | *LV1* | *-0.61 [-0.69 -0.53]* | ***<0.001*** |
| | *LV2* | *-0.19 [-0.26 -0.11]* | ***<0.001*** |
| *Lexical Fluency* | *LV1* | *-0.52 [-0.61 -0.44]* | ***<0.001*** |
| | *LV2* | *-0.20 [-0.28 -0.12]* | ***<0.001*** |
| *Memory* | *LV1* | *-0.42 [-0.51 -0.33]* | ***<0.001*** |
| | *LV2* | *-0.13 [-0.22 -0.04]* | ***0.003*** |
| *Inductive Reasoning* | *LV1* | *-0.82 [-0.88 -0.77]* | ***<0.001*** |
| | *LV2* | *0.13 [0.08 0.19]* | ***<0.001*** |
| *Mathematical Reasoning* | *LV1* | *-0.82 [-0.88 -0.76]* | ***<0.001*** |
| | *LV2* | *0.13 [0.08 0.19]* | ***<0.001*** |
| *Verbal Reasoning* | *LV1* | *-0.78 [-0.84 -0.71]* | ***<0.001*** |
| | *LV2* | *0.12 [0.07 0.18]* | ***<0.001*** |

In section 2.4 of the main text we present results of linear models conducted using square root and inverse normalized data to account for count nature and skew. Here we present results of linear models of the same form, but using untransformed data for comparison (Table S3). For each model, significance of the terms of interest does not change.

**Table S3:** Statistical results from linear models run for Wave 12 performance on each of 6 cognitive tests, using untransformed data. Each model had the form $Test \sim LV1 + LV2 + Age + Sex + Education.$ Unstandardized beta coefficients and 95% confidence intervals are reported. Bolded p values are deemed significant at a bonferroni threshold of p<0.0083.

| Test | Term | β [95% CI] | p |
|---|---|---|---|
| Semantic Fluency | LV1 | -0.84 [-0.95 -0.73] | **<0.001** |
| | LV2 | -0.48 [-0.68 -0.29] | **<0.001** |
| Lexical Fluency | LV1 | -0.84 [-1 -0.68] | **<0.001** |
| | LV2 | -0.63 [-0.92 -0.34] | **<0.001** |
| Memory | LV1 | -0.36 [-0.44 -0.28] | **<0.001** |
| | LV2 | -0.22 [-0.37 -0.08] | **0.0024** |
| Inductive Reasoning | LV1 | -3.35 [-3.56 -3.13] | **<0.001** |
| | LV2 | 1.13 [0.74 1.51] | **<0.001** |
| Mathematical Reasoning | LV1 | -1.84 [-1.96 -1.72] | **<0.001** |
| | LV2 | 0.62 [0.39 0.85] | **<0.001** |
| Verbal Reasoning | LV1 | -1.5 [-1.62 -1.39] | **<0.001** |
| | LV2 | 0.51 [0.3 0.72] | **<0.001** |
